# Control of Golgi Architecture Through Three Centrosome Cohesion Pathways

**DOI:** 10.64898/2026.10.15.759098

**Authors:** Aadyasha Mishra, Sietse J. Dijt, Tobias Benedikt Roth, Laura Theile, Annett Neuner, Simon Anders, Rifka Vlijm, Elmar Schiebel

## Abstract

Centrosome cohesion mechanisms link the two interphase centrosomes into a single microtubule-organizing center that is closely associated with the Golgi apparatus. The contribution of cohesion mechanisms to Golgi organization and function remains unclear. Here, we show that rootletin-CEP68 filaments, the C-Nap1-ninein linker, and microtubules derived from subdistal appendages, all of which promote centrosome cohesion, regulate the Golgi through distinct mechanisms while acting synergistically to maintain its organization and function. Centrosome-anchored rootletin-CEP68 filaments associate with AKAP450 through periodic binding of its PACT domain to CEP68 and extend into the Golgi, thereby establishing a physical connection between both organelles. The rootletin-CEP68-AKAP450 linkage even functions in cell types in which rootletin and AKAP450 do not mediate centrosome cohesion, suggesting that Golgi organization is the basic function of these filaments. Knockout analyses demonstrate that the three centrosome cohesion mechanisms collectively regulate Golgi morphology, positioning, and reassembly as well as cell migration. Each mechanism contributes distinctly to the overall process, indicating that they operate as a multifaceted system.

## Introduction

The centrosome is the primary microtubule-organizing center of animal cells and is essential for the cell organization as well as accurate chromosome segregation during mitosis. Each centrosome consists of centrioles, which are cylindrical microtubule triplet-based structures, approximately 250 nm in diameter and 500 nm in length, and the pericentriolar material (PCM) that surrounds the centrioles (EA Nigg, 2011). While the centriole contributes structural stability to maintain centrosome integrity, the PCM recruits the kinase PLK4, an essential regulator of centriole duplication during S phase and contains with pericentrin (PCNT) and CDK5RAP2 (CDK5 regulatory subunit-associated protein 2) factors that regulate microtubule nucleation by the γ-tubulin ring complex (EA Nigg, 2011; Liu *et al.,* 2019; Song *et al*., 2026).

Daughter centrioles, assembled during S phase, remain attached to the mother centriole and gradually mature into functional centrosomes by recruiting centrosomal proteins from G2 phase through mitosis (Izquierdo *et al*., 2014). Distal and subdistal appendages are protein assemblies located at the distal end of mother centrioles. They are absent from daughter centrioles until two rounds of mitosis have occurred following their assembly. Subdistal appendages contain a number of proteins including CEP128 and ninein that anchor stable microtubules to the centrosome during interphase (Hall & Hehnly, 2021; Kashihara *et al*., 2019; Mönnich *et al*., 2018). After duplication, the perpendicularly arranged mother and daughter centrioles disengage at the end of mitosis (Tsou & Stearns, 2006). Subsequently, the two disengaged centrosomes become connected by the centrosome linker filaments that attach to the proximal end of the two centrioles via the protein C-Nap1 (centrosomal Nek2-associated protein 1) (Dang & Schiebel, 2022; Fry *et al.,* 1998; Mayor *et al*., 2000). C-Nap1 organizes rootletin–CEP68 filaments that mediate centrosome cohesion in U2OS (human osteosarcoma) and HCT116 (human colorectal carcinoma) cells. Within these filaments, rootletin molecules are arranged in a polar fashion at defined intervals, with N-to-N and C-to-C spacing of 75 nm. CEP68 binds to rootletin via its C-terminal spectrin motif and crosslinks rootletin fibers into bundles (Vlijm *et al.,* 2018). In addition, C-Nap1 interacts with ninein at the proximal end of centrioles, contributing to centrosome cohesion during interphase through an unclear mechanism (Theile *et al.,* 2023). Interestingly, C-Nap1 and ninein also impact microtubule nucleation, as evidenced by delayed microtubule regrowth from centrosomes in knockout (KO) cell lines in a depolymerization/polymerization assay. Reduced recruitment of γ-tubulin to the centrosomes may partly explain this phenotype (Theile *et al.,* 2023). The C-Nap1–ninein pathway is required for centrosome cohesion in all tested cell lines, functionally overlapping with the rootletin-CEP68 filaments in HCT116 and U2OS cells. Interestingly, although rootletin-CEP68 filaments are formed in RPE1 (human retinal pigment epithelial) cells with the same periodicity as in U2OS and HCT116 cells (Vlijm *et al.,* 2018), they do not contribute to centrosome cohesion (Theile *et al.,* 2023). This raises the question of whether rootletin–CEP68 filaments have additional functions at centrioles.

In addition to the centrosome linker, microtubules anchored to the subdistal appendages of the mother centriole and to the PCM of the daughter help pulling the two centrosomes together through the activity of the minus-end-directed, tetrameric motor protein KIFC3 (Hata *et al.,* 2019). Consequently, even without the centrosome linker, centrosomes stay paired through microtubule- and KIFC3-mediated interactions, although this linkage is more dynamic than the one maintained by the centrosome linker (Hata *et al.,* 2019). Simultaneous disruption of the distal end microtubule pathway and the proximal end cohesion mechanisms leads to nearly complete centrosome disjunction in interphase cells (Theile *et al.,* 2023).

AKAP450 (protein kinase A-anchoring protein; also known as AKAP9, AKAP350 or CG-NAP) localizes to both the centrosome and the Golgi apparatus (Rivero *et al.,* 2009). Together with pericentrin, AKAP450 contains a C-terminal pericentrin-AKAP450 centrosome-targeting (PACT) domain that is sufficient to recruit proteins to the PCM (Gillingham & Munro, 2000). Besides PCM localization, AKAP450 has also been described as a component of the proximal-end filaments that interconnect the two centrioles, consistent with its interaction with CEP68 (Kolobova *et al.,* 2017); however, the functional significance of this AKAP450 pool remains unclear.

AKAP450 is anchored to the Golgi through its interaction with the cis-Golgi protein GM130 (Rivero *et al.,* 2009). In microtubule regrowth assays, Golgi-associated AKAP450 is required for microtubule nucleation at the Golgi, whereas centrosomal microtubule nucleation appears unaffected by AKAP450 depletion (Rivero *et al.,* 2009; Wu *et al.,* 2016).

From previous work, it is clear that the Golgi apparatus and the centrosome are dynamically linked during interphase (Sütterlin & Colanzi, 2010). Positioning of the centrosome–Golgi complex toward the leading edge of a migrating cell is important for directed cell migration because it helps organize polarized, microtubule-dependent trafficking of vesicles toward the leading edge (Mascanzoni *et al.,* 2024). However, how the centrosome and Golgi interact to establish and maintain this dynamic and polarized organization remains incompletely understood.

Here, we analyzed how three centrosome-derived elements, C-Nap1-ninein *(Theile et al.,* 2023), the microtubule centrosome cohesion pathway (Hata *et al.,* 2019), and the rootletin-CEP68 filaments (Bahe *et al.,* 2005, Graser *et al.,* 2007), that have the potential to function in centrosome cohesion, organize the Golgi apparatus during interphase. Rootletin-CEP68 filaments are particularly interesting because, despite forming similarly to those described in cell lines in which they function as centrosome linkers, they do not contribute to centrosome cohesion in RPE1 cells, raising the question about their general function (Theile *et al*., 2023). We found that AKAP450, by binding to CEP68 through its PACT domain, physically connects the rootletin-CEP68 filaments to the Golgi. This connection, together with the C-Nap1 centrosome linker and the microtubule pathway, determines the compactness of Golgi stacks and their reformation after dissolution by the drug brefeldin A. Additionally, each of these three cohesion mechanisms influences cell behavior, including cell migration, in a distinct manner. Our study identifies a previously unrecognized organizational role for centrosome-associated rootletin–CEP68 fibers as Golgi organizers, independent of their established function in centrosome cohesion. This function likely reflects the conserved cellular role of rootletin–CEP68 filaments.

## Results

### AKAP450 decorates rootletin fibers extending from the centrosome

AKAP450 is a large, multidomain protein containing several coiled-coil regions, an N-terminal GM130-binding site, and a C-terminal PACT domain (Fig 1A). It localizes to both the centrosomes and the Golgi apparatus (Rivero *et al*., 2009; Kolobova *et al*., 2017). To determine whether the centrosomal localization of the protein is cell type dependent, we examined three distinct cell lines - RPE1, HCT116, and U2OS - during both interphase and mitosis. In all three cell lines, AKAP450 localized to the centrosome during interphase (Fig 1B). It was predominantly positioned between the two γ-tubulin signals marking the centrioles (Fig 1C; Appendix Fig S1A and B). Quantification of the AKAP450 signal intensity at centrosomes during interphase indicated a 1.5-fold higher intensity in U2OS and HCT116 cells compared to RPE1 cells (Fig 1D). This localization pattern is consistent with AKAP450 residing at the centrosome linker in interphase cells.

**Figure 1.**
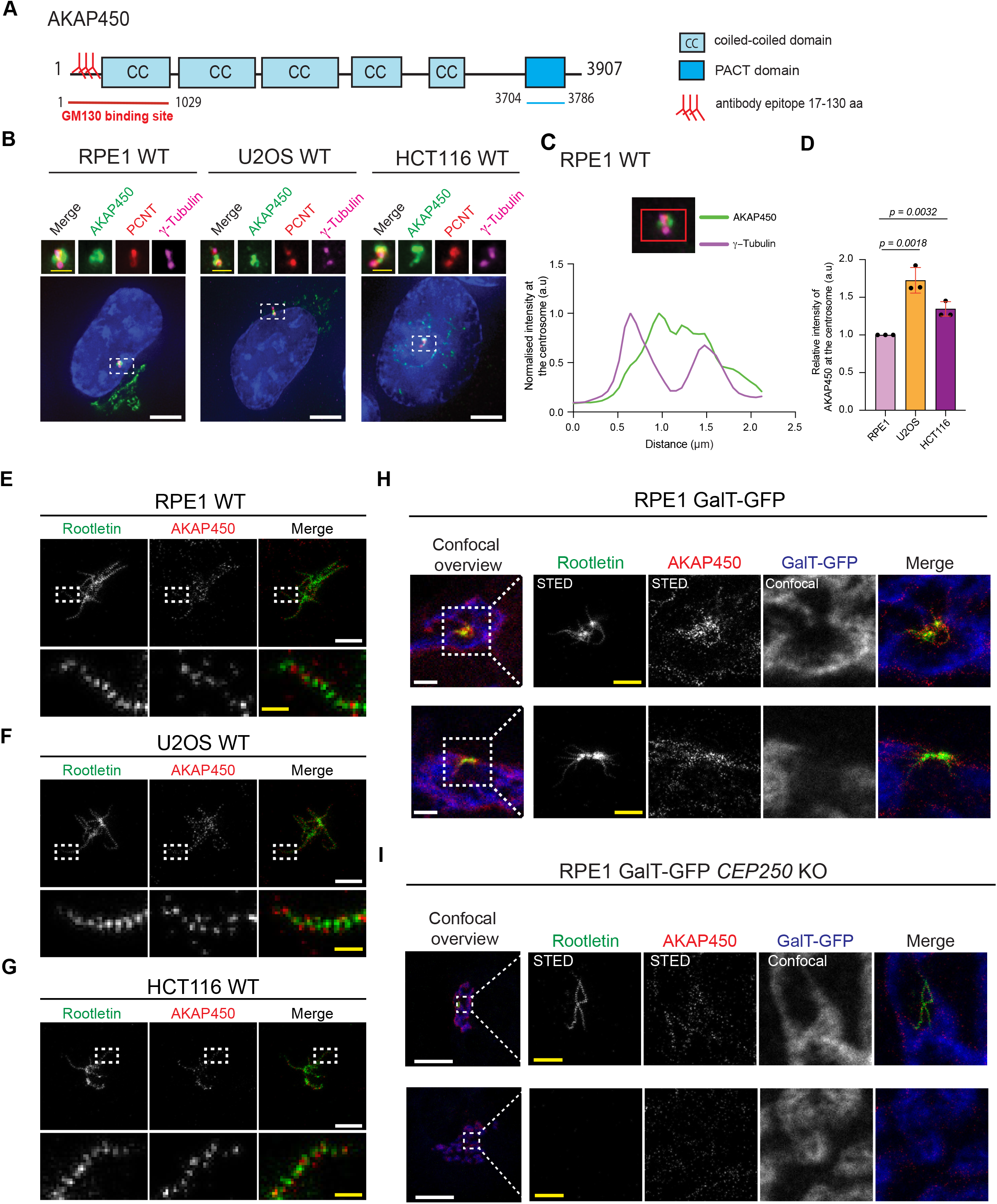
Association of periodic rootletin-AKAP450 filaments with the Golgi apparatus. **A** Domain organization of AKAP450 consisting of multiple coiled-coil domains and the PACT domain at the C-terminus. The antibody epitope is at the N-terminus between 17-130 aa (indicated by Y). The GM130 binding site and the PACT domain are indicated. **B** Representative fluorescent images of AKAP450 (green) at the centrosomes and Golgi in wild type RPE1, HCT116 and U2OS cells. γ-tubulin (magenta) and PCNT (red) were used to mark centrosomes and DNA was stained with DAPI. The enlargements on top show the centrosome signals of the boxed area in the main panel. White scale bars, 5 µm. Yellow scale bars, 1 µm. **C** Representative line-scan intensity profile of a centrosome (centrosome shown on top; from cell in (B), left) showing the centrosomal distribution of AKAP450 (green) and γ-tubulin (magenta) in RPE1 cells. Fluorescence intensity (arbitrary units, a.u.) is plotted as a function of distance (µm) along the x axis. n=15 cells from 3 independent experiments were analyzed. **D** Quantification of the signal intensity of AKAP450 at the centrosome (a.u) in (B). n >100 cells per experiment, N=3 independent experiments. Bar and error represent mean and SD. Unpaired t test. **E-G** STED images of AKAP450 (red) and rootletin (green) at centrosomes in RPE1 (E), U2OS (F), and HCT116 (G) cells. For each image, the area indicated by the white rectangle is shown enlarged below the image to show the typical organization of AKAP450 along rootletin fibers. White scale bars, 1 µm. Yellow scale bars, 200 nm. **H** STED images of AKAP450 (red), rootletin (green), and confocal images of GalT-GFP (blue) at centrosomes in RPE1 GalT-GFP WT cells, showing examples of rootletin fibers going along or towards GalT-GFP. Note, one image from (H) is also shown in Appendix Fig. S2C. White scale bars, 2 µm. Yellow scale bars, 1 µm. **I** STED images of AKAP450 (red), rootletin (green), and confocal images of GalT-GFP (blue) at centrosomes in RPE1 GalT-GFP *CEP250* KO cells, showing that there are detached rootletin fibers at the Golgi. White scale bars, 10 µm. Yellow scale bars, 1 µm.

We next examined AKAP450 localization during mitosis, as disassembly of the centrosome linker during late G2/M is expected to reduce the centrosomal AKAP450 signal if the centrosome linker is its primary binding site. In all three cell lines, AKAP450 association with the centrosome was 12 times (RPE1 and U2OS) or 5 times (HCT116) diminished during mitosis (Appendix Fig S1C-F). As in interphase cells, the mitotic centrosomal AKAP450 signal was reduced 1.7-3 fold in RPE1 cells compared with HCT116 and U2OS cells (Appendix Fig S1F).

Previously, we analyzed the organization of rootletin-CEP68 linker filaments using STED super-resolution microscopy, illustrating their periodic arrangement (Vlijm *et al*., 2018). In RPE1, U2OS and HCT116 WT cells, rootletin and AKAP450 both form extended, fiber-like structures radiating from the centrosome, visible by STED imaging. In magnified views (lower panels), rootletin and AKAP450 appear as discrete puncta. The AKAP450 puncta are positioned around the rootletin fibers, rather than as a continuous co-localized signal (Fig 1E-G; Appendix Fig. S2A). In addition, AKAP450 was not enriched at rootletin intersections corresponding to γ-tubulin-positive centrioles (Appendix Fig. S2A), consistent with our immunofluorescence analysis, which identified centrioles as a minor, rather than a major, site of AKAP450 localization (Fig. 1B). In addition, in 6–7% of RPE1, U2OS, and HCT116 cells, rootletin filaments were detached from the centrosomes. Notably, these detached rootletin filaments were also decorated with AKAP450 (Appendix Fig. S2B).

Interestingly, in RPE1 cells the rootletin-AKAP450 filaments ended in Golgi areas marked by the Golgi marker GalT-GFP (imaged in confocal mode) (Fig 1H; Appendix Fig. 2C and D), indicating contacts between rootletin filaments and the Golgi. In the majority of RPE1 cells (83%), the rootletin–AKAP450 filaments were detected as a web-like network connecting the GalT-GFP–marked Golgi area (Fig 1E, Appendix Fig S2C). In *CEP250* knockout (KO) cells (coding for C-Nap1), in which rootletin-AKAP450 filaments are no longer anchored to the centrosome, 73% of cells contained detached rootletin filaments within the cytoplasm. Among these cells, 98% (72% of *CEP250* KO cells) showed rootletin-AKAP450 filaments at the GalT-GFP-marked Golgi apparatus (Fig 1I, Appendix Fig S2E and F). Only 1% of *CEP250* KO cells contained rootletin-AKAP450 filaments that were not associated with the Golgi (Appendix Fig S2E). 27% of *CEP250* KO cells did not contain rootletin filaments. Together, these results indicate that AKAP450 decorates rootletin filaments that extend from the proximal end of the two centrioles into Golgi regions, where it likely interacts with its binding partner GM130 (Rivero *et al*., 2009). In contrast, AKAP450 exhibits no or only weak PCM localization.

**Figure 2.**
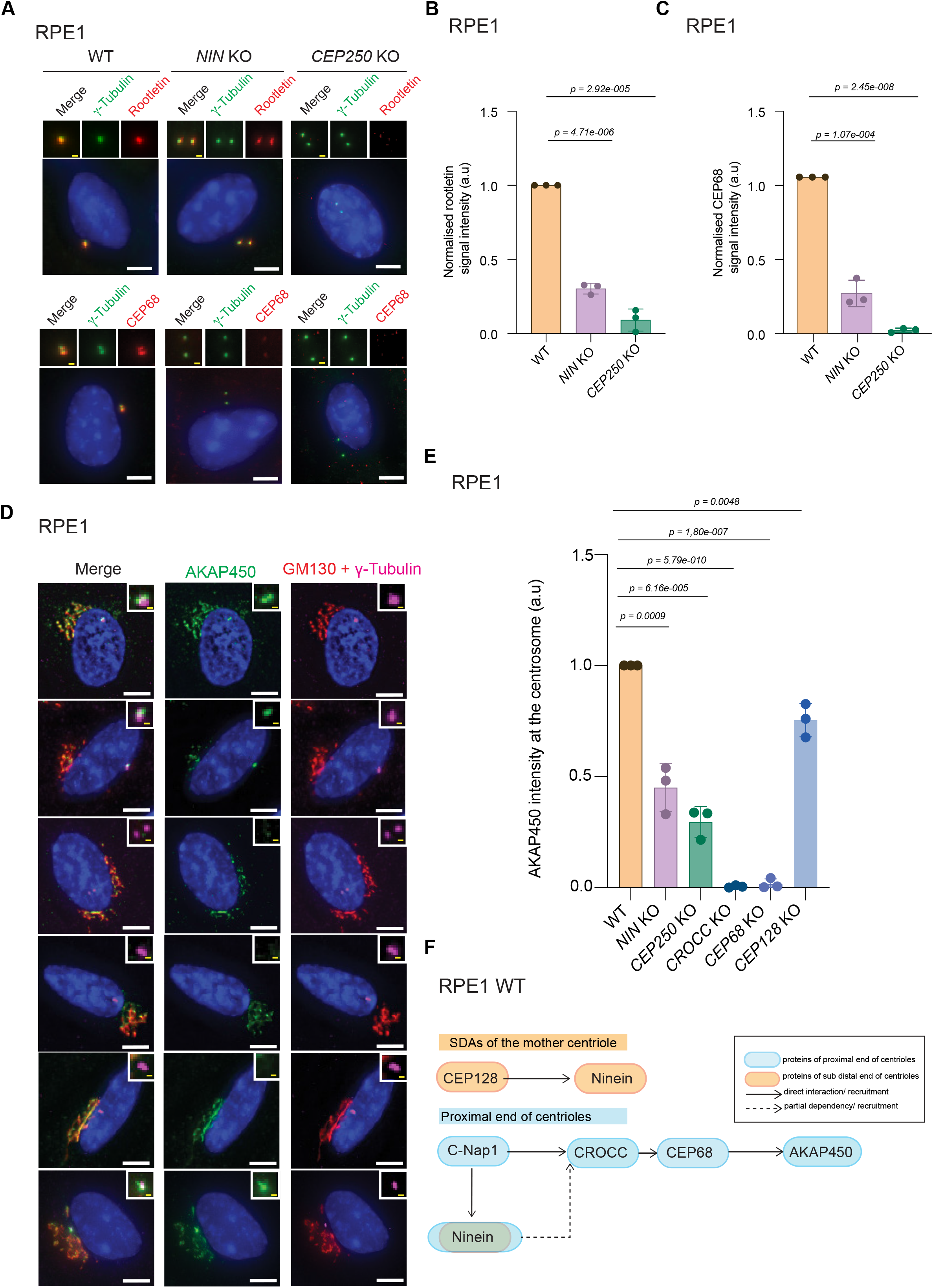
Impact of the centrosome linker on AKAP450 localizations with centrosomes. **A** Representative fluorescent image of RPE1 WT, *NIN* KO, *CEP250* KO cells. Rootletin (red) and CEP68 (red) were stained. γ-tubulin (green) was used to mark centrosomes and DNA was stained with DAPI. The enlargements at the top show the centrosome signal of the cell below. White scale bars, 5 µm. Yellow scale bars, 1 µm. **B, C** Quantification of the normalized CROCC (B) and CEP68 (C) signal intensities (a.u.) at the centrosome in (A). (B) n>110, (C) n>105 cells per experiment, N=3 independent experiments. Bar and error represent mean and SD. Unpaired *t*-test. **D** Representative fluorescent image of AKAP450 (green) at the centrosomes and Golgi in RPE1 WT, *NIN* KO, *CEP250* KO, *CROCC* KO, *CEP68* KO, *CEP128* KO cells. γ-tubulin (magenta) was used to mark centrosomes, GM130 (red) was used to mark the cis face of the Golgi and DNA was stained with DAPI. The enlargements at the top show the centrosome signal of the cell in the main panel. White scale bars, 5 µm. Yellow scale bars, 1 µm. **E** Quantification of the normalized AKAP450 signal intensity (a.u.) at the centrosome in (D). n>100 per experiment, N=3 independent experiments. Bar and error bars represent mean and SD. Unpaired *t*-test. **F** Summary of centrosome linker localizations and functions at the centrosome, based on the results from (A), (B), (C) and (D) and Fig EV2.

### Impact of the centrosome linker on AKAP450 localizations with centrosomes

To investigate the role of centrosome cohesion elements in AKAP450 recruitment to centrosomes, we analyzed AKAP450 association in published *CEP250* (coding for C-Nap1), *CEP68*, *CROCC* (coding for rootletin) and *NIN* (coding for ninein) RPE1 cell lines (Theile *et al*., 2023). To affect MTs organized by subdistal appendages, we constructed *CEP128* KO, *CEP128*-*250* KO and *CEP128*-*68* KO cells as outlined in Appendix Fig S3A-C. The *CEP128* KO cell lines lacked an CEP128 signal by immunoblotting and a centrosomal CEP128 signal by indirect immunofluorescence (Appendix Fig S3D-F). Centrosome cohesion was disrupted in *CEP128-250* KO cells, whereas it remained intact in *CEP128* KO and *CEP128-68* KO cells, owing to the presence of the C-Nap1-ninein linker (Appendix Fig S3E and G). We also examined centrosomes in *CEP128* KO cells by electron microscopy and compared their ultrastructure with that of centrosomes in *NIN* KO cells. *CEP128* KO centrioles lacked any detectable subdistal appendages (Appendix Fig S3H), whereas *NIN* KO centrioles retained electron-dense structures at the distal end of the centrioles, corresponding to subdistal appendages (Fig EV1A). These observations are consistent with CEP128 acting upstream of ninein in the hierarchy of subdistal appendage assembly (Hall & Hehnly, 2021).

Using these cell lines, we first determined the hierarchical organization of centrosome linker components in RPE1 cells, which had previously only been determined in HCT116 cells (Theile *et al*., 2023). Interestingly, unlike in HCT116 cells, where centrosomal localization of rootletin/CEP68 is independent of ninein (Theile *et al*., 2023), deletion of *NIN* in RPE1 cells reduced the centrosomal levels of rootletin and CEP68 by ca. 70% (Fig 2A-C), while lack of *CROCC* (rootletin) or *CEP68* did not affect ninein signal intensity at centrosomes (Fig EV1B and C). Lack of C-Nap1 (*CEP250* KO) reduced ninein levels by only 50%, likely due to the pool of ninein localized at subdistal appendages. This reduction was further increased in C*EP128-250* double KO cells (Fig EV1C) (Hall & Hehnly, 2021). Together, these findings suggest that, in RPE1 cells, rootletin/CEP68 filaments can be recruited to centrosomes through two distinct mechanisms: one mediated by C-Nap1 and the other by ninein.

We next examined how AKAP450 depends on centrosomal linker proteins in RPE1 cells. In agreement with the STED data (Fig 1), AKAP450 association with centrosomes was fully dependent on rootletin (*CROCC* KO) and CEP68 (*CEP68* KO) (Fig 2D and E). Consistent with the partial dependence of rootletin and CEP68 on ninein (Fig 2A-C), centrosomal localization of AKAP450 was also reduced in *NIN* KO cells (Fig 2D and E). The reduction in rootletin and CEP68 signals at the centrosomes of *NIN* KO cells was rescued by Tet_ON_-induced expression of *NIN* in *NIN* KO cells (compare Fig EV1D-F with Fig. 2D and E). Although the centrosomal localization of rootletin and CEP68 depends on C-Nap1 and is required for the recruitment of AKAP450 to centrosomes, a weak centrosomal AKAP450 signal persisted in *CEP250*-deficient cells (Fig 2D and E). Since AKAP450 can associate with the PCM (Keryer *et al*., 2003), it is plausible that loss of C-Nap1 exposes a PCM-binding site for AKAP450 at the centrosome.

Finally, we investigated how loss of AKAP450 affects the rootletin and CEP68 in RPE1 cells. Previous work indicated that siRNA-mediated depletion of AKAP450 leads to the loss of CEP68 from centrosomes in U2OS cells (Kolobova *et al*., 2017). Our data show that knockout of *AKAP450* in RPE1 cells did not affect the localization of CEP68 or rootletin at centrosomes (Fig EV1G-I). These results suggest that AKAP450 is recruited to centrosomes via rootletin-CEP68 filaments without affecting their centrosomal recruitment. A summary of these localization interdependency data is shown Fig 2F.

### The PACT domain of AKAP450 interacts with the centrosome linker protein CEP68

Previous studies reported that CEP68 can bind to AKAP450 (Graser *et al*.,2007; Kolobova *et al*., 2017; Vlijm *et al*., 2018; Wu *et al.,* 2016). We sought to identify the region of AKAP450 responsible for mediating this interaction using the constructs shown in Fig 3A. Immunoprecipitation experiments revealed that the C-terminal AKAP450 fragment spanning amino acids 3000–3907, which contains the PACT domain, interacted with CEP68, whereas the N-terminal region (aa 1–1029) did not (Fig 3B). Notably, the PACT domain of AKAP450 alone (aa 3704-3907) was sufficient to immunoprecipitate CEP68 (Fig 3B, lane 7). We established that AKAP450^3704-3907^ did not bind to the HA beads without *CEP68-HA* expression (Fig 3B, lane 3).

**Figure 3.**
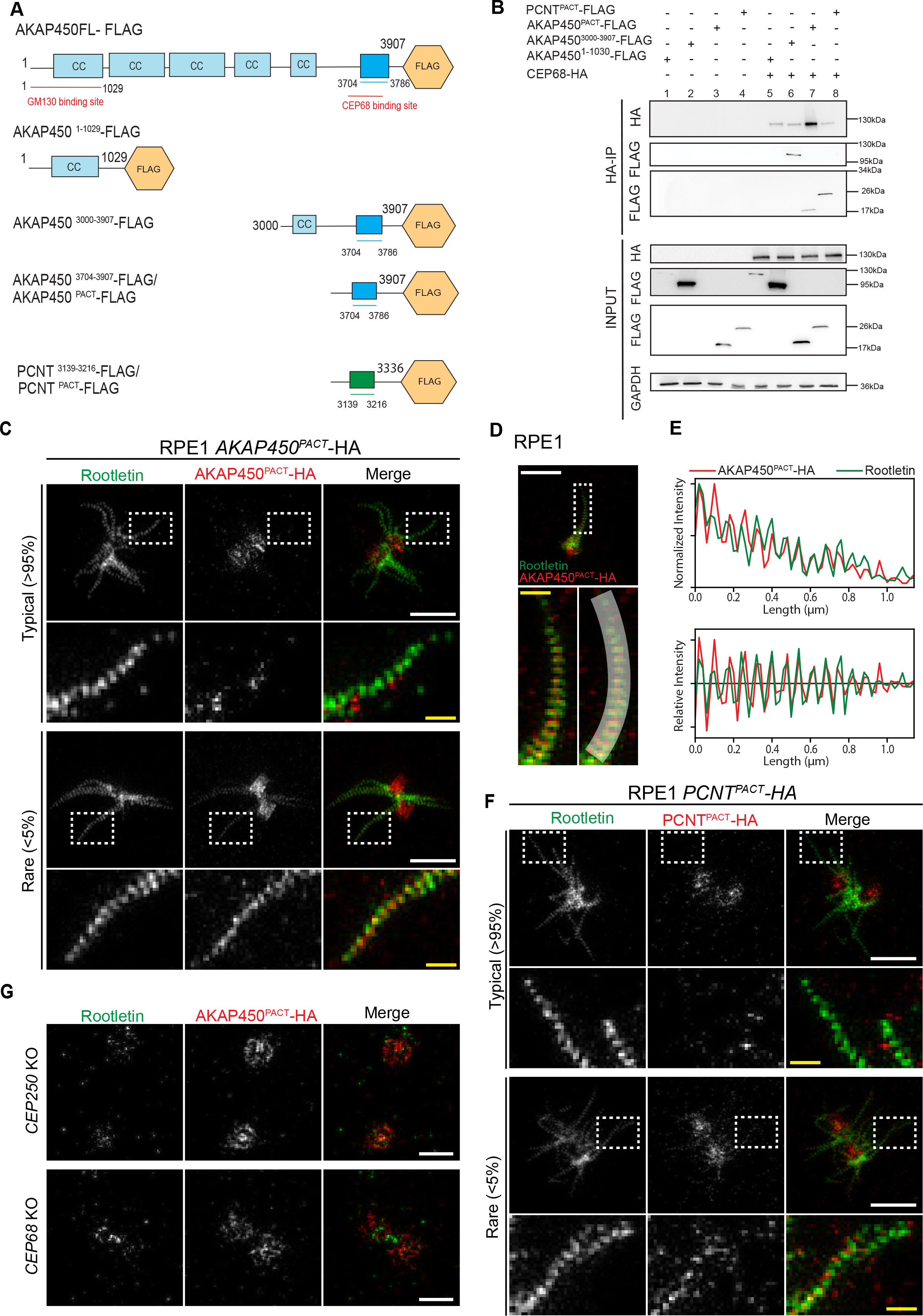
PACT domain is necessary for AKAP450 to localise to the centrosome linker via CEP68. **A** Summary of FLAG-tagged constructs of AKAP450 and PCNT used for immunoprecipitation (IP) experiments. **B** Representative HA IP from HEK293T cells expressing HA-tagged CEP68 and FLAG-tagged subdomains of AKAP450 and PCNT. GAPDH was used as an input control. N=2 biologically independent experiments with equal outcome. **C** STED images of AKAP450^PACT^-HA (red) and rootletin (green) at centrosomes in RPE1 AKAP450^PACT^-HA cells. Typically, there is little AKAP450^PACT^-HA around rootletin fibers, but in rare cases, the AKAP450^PACT^-HA signal clearly follows the rootletin fibers. The regions indicated by the white rectangles are shown enlarged below the images. White scale bars, 1 µm. Yellow scale bars, 200 nm. **D** STED image of AKAP450^PACT^-HA (red) and rootletin (green) with a line drawn outward from the centriole along the fiber that is indicated in the enlarged image. White scale bars, 1 µm. Yellow scale bars, 200 nm. **E** Intensity profile along the line drawn in (D), showing the normalized intensity (top panel) and relative intensity compared to a 3-pixel moving average (bottom panel). The profile shows AKAP450^PACT^-HA intensity closely follows the periodic rootletin signal. **F** STED images of PCNT^PACT^-HA (red) and rootletin (green) at centrosomes in RPE1 PCNT^PACT^-HA cells, showing the organization of PCNT^PACT^-HA around rootletin fibers is similar to AKAP450^PACT^-HA. White scale bars, 1 µm. Yellow scale bars, 200 nm. **G** Typical STED images of AKAP450^PACT^-HA (red) and rootletin (green) at the centrosome in RPE1 *CEP250* KO and *CEP68* KO cells with AKAP450^PACT^-HA overexpression. White scale bars, 500 nm.

The PACT domains of AKAP450 and PCNT are highly conserved (Gillingham & Munro, 2000), suggesting that PCNT^PACT^ may also be capable of interacting with CEP68. To test this hypothesis, we performed co-immunoprecipitation experiments and detected an interaction between CEP68 and PCNT^PACT^ (Fig 3A and B, lane 8; control lane 4), indicating that the PACT domains of both AKAP450 and PCNT are sufficient to mediate interaction with CEP68.

Based on their ability to bind CEP68 and recruit fusion proteins to the PCM (Gillingham & Munro, 2000), we hypothesized that the PACT domains of AKAP450 (AKAP450^PACT^) and PCNT (PCNT^PACT^) would exhibit dual localization to both the PCM and the centrosome linker. To test this, we examined the subcellular distribution of the PACT domains by STED microscopy. Doxycycline-inducible AKAP450^PACT^-HA and PCNT^PACT^-HA constructs were expressed in RPE1 WT cells. STED imaging of RPE1 WT cells revealed that both PACT fragments localized to the centrosomal PCM/centrioles, as indicated by the HA signal at the intersection of rootletin filaments corresponding to the γ-tubulin-positive centrioles (Fig 3C and F - top, Appendix Fig S4A and B). AKAP450^PACT^ signal was also detected sparsely along rootletin filaments (Fig 3C and D). Complete decoration of the filaments was with 1-5% of the cases relatively rare (Fig 3C, bottom, Fig 3D, Appendix Fig S5A, bottom). In instances of complete decoration, AKAP450^PACT^ followed the periodicity of the rootletin filaments (Fig 3D and E).

To determine the requirements for AKAP450^PACT^ domain binding to centrosomes, we used RPE1 *CEP250* KO and *CEP68* KO cell lines. In *CEP250* KO cells, AKAP450^PACT^ was associated with a barrel-like structure, most likely corresponding to the outer surface of the centriole wall, as well as a hollow region surrounding the centrioles (Fig 3G, top, Appendix Fig S5B). Centrosome-detached rootletin filaments in *CEP250* KO cells were not decorated by AKAP450^PACT^ (Appendix Fig S5C), further supporting the low affinity of AKAP450^PACT^ for rootletin filaments. In RPE1 *CEP68* KO cells, that contain short rootletin filaments (Vlijm *et al*., 2018), the AKAP450^PACT^ localized to centrioles and the hallow but not the remaining rootletin filaments (Fig 3G, bottom, Appendix Fig S5D) indicating that CEP68 is essential for PACT recruitment to rootletin filaments. Similar observations were made for PCNT^PACT^ (Fig 3F, Appendix Fig S6A-D).

These data indicate that the PACT domain preferentially binds to centrioles, whereas binding to rootletin-CEP68 filaments was inefficient. Given that AKAP450 predominantly associates with rootletin-CEP68 filaments, these findings suggest that AKAP450 employs mechanisms to suppress the binding of its PACT domain to centrioles, thereby favoring its association with rootletin–CEP68 filaments.

### AKAP450 functions as a centrosome linker component in U2OS and HCT116 cells, but not in RPE1 cells

The localization of AKAP450 to rootletin/CEP68 filaments suggests that it may play a functional role in centrosome cohesion. To test this, we depleted C-Nap1, rootletin and AKAP450 using siRNA in RPE1, U2OS, and HCT116 cells (Fig EV2). The depletion efficiency was in the range from 70%-97% as determined by the reduction of the centrosomal signal (Appendix Fig S7). In addition, centrosomal microtubule cohesion was disrupted by treatment with the drug nocodazole (Fig EV2). In RPE1 cells, where centrosome cohesion is primarily maintained by the C-Nap1-ninein pathway and not by rootletin-CEP68 (Theile *et al*., 2023), AKAP450 depletion did not affect centrosome cohesion, even after microtubule depolymerization with nocodazole (Fig EV2A and B). In contrast, in HCT116 and U2OS cells, where centrosome cohesion depends on both the rootletin-CEP68 and C-Nap1-ninein pathways, loss of AKAP450 impaired centrosome cohesion particularly when the microtubule pathway was disrupted by the addition of nocodazole (Fig EV2C-F). These findings indicate that AKAP450 contributes to centrosome linker function in U2OS and HCT116 cells but not in RPE1 cells.

To validate the findings obtained from fixed-cell analyses, we performed live-cell imaging of RPE1 and HCT116 cells expressing the centrosomal marker γ-tubulin–mRuby2. Cells were transfected with either a non-specific control siRNA (siControl) or an AKAP450-targeting siRNA (Fig 4, Movie EV1, Movie EV2). In wild-type RPE1 cells, microtubule depolymerization by nocodazole had only a mild little effect on centrosome cohesion. Among the 15 cells analyzed, only two exhibited a transient separation of the two centrosomes (Fig 4B, purple and blue lines). The cell indicated by the purple line rejoined the two centrosomes after approximately 50 min. In the remaining cells, the two centrosomes remained associated (< 2 µm) throughout the imaging period. A similar behavior of RPE1 was observed following AKAP450 depletion with and without nocodazole addition (Fig 4C and D, Movie EV1) indicating that AKAP450 does not have a centrosome linker function in RPE1 cells. In contrast, depletion of AKAP450 partially relieved centrosome cohesion in HCT116 cells, as indicated by the strong separation of the two centrosomes overtime (compare Fig 4E and F, control with Fig 4G and H, siAKAP450, Movie EV2). Addition of nocodazole further enhanced this defect in HCT116 cells as the average distance of centrosome separation increased from 3 µm to 6 µm (Fig 4H). Together, these findings demonstrate that the contribution of AKAP450 to centrosome cohesion is cell type-specific and mirrors the requirement for rootletin (Theile *et al*., 2023).

**Figure 4.**
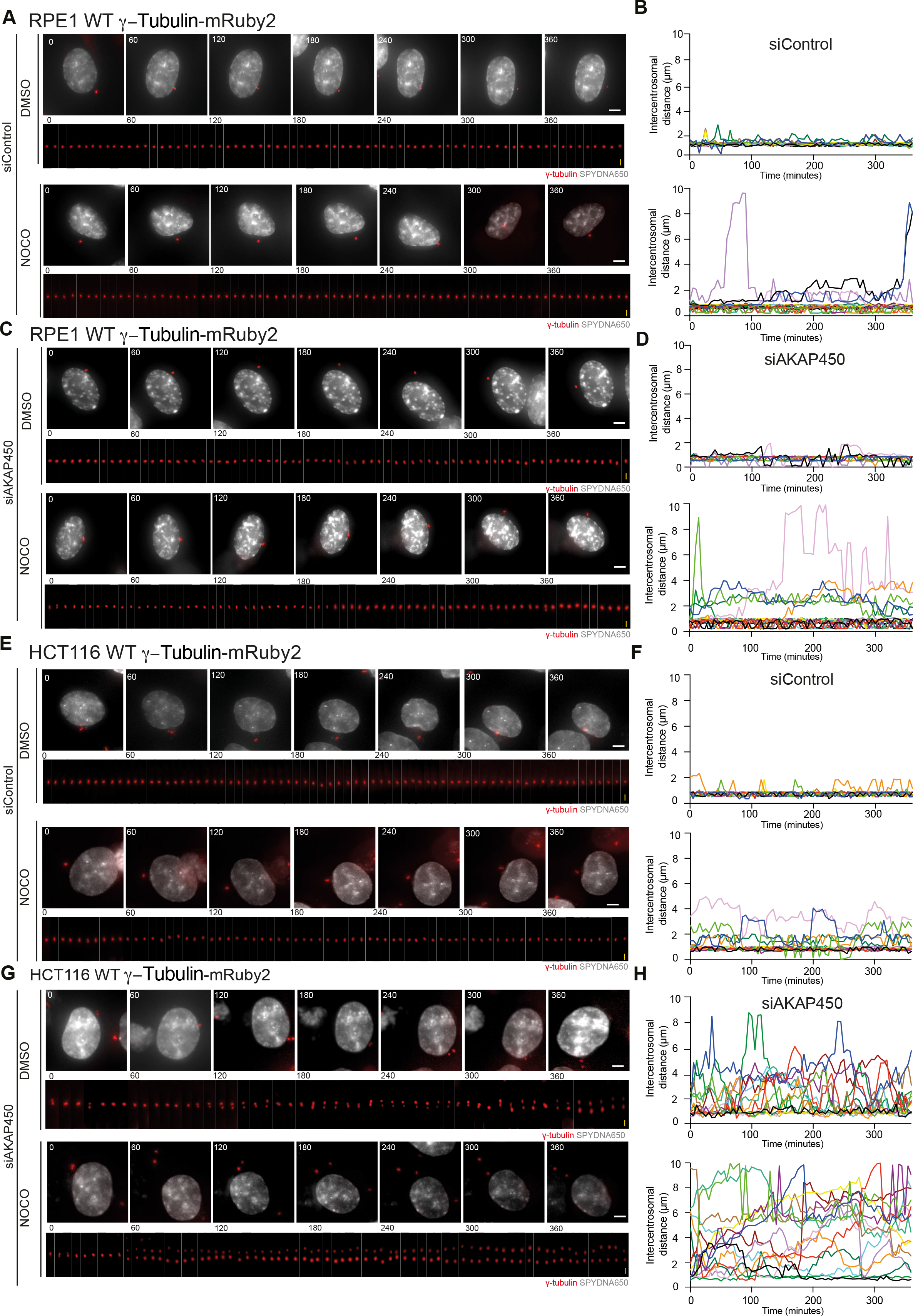
AKAP450 is a component of the centrosome linker in HCT116 and U2OS cells. **A, C, E, G** Time-lapse images (5 min intervals) of RPE1 WT and HCT116 WT cells expressing γ-tubulin-mRuby2 (red) as markers for the centrosomes. Cells were under siRNA depletion of non-specific siRNA control (siControl) and AKAP450 siRNA, with and without Nocodazole (NOCO) treatment. Representative still images (top rows) and kymographs of centrosomes (bottom rows) are shown. The DNA was marked with SPY DNA650 (grey). White scale bars, 5 µm. Yellow scale bars (vertical), 2 µm. (Movie EV1, EV2). **B, D, F, H** Intercentrosomal distances (µm) from (A), (C), (E) and (G) are plotted over time (n=15 cells, pooled from N=3 independent experiments).

### Centrosome cohesion factors connect the centrosome to the Golgi

The dual localization of AKAP450 at both the centrosome linker and the centrosome, together with the overlap of rootletin–CEP68–AKAP450 filaments with the Golgi (Fig. 1), suggests that these filaments physically connect the Golgi apparatus to centrosomes. Furthermore, rootletin filaments, in concert with centrosome cohesion mechanisms, the C-Nap1–ninein linker and subdistal appendage-derived microtubules (Hata *et al*., 2019; Theile *et al*., 2023), may contribute to Golgi organization by regulating the distance between the two centrosomes and, consequently, the spatial distribution of rootletin filaments throughout the cell.

To test this hypothesis, we analyzed the centrosome–Golgi connection in fixed RPE1 cells with defects in centrosome cohesion and rootletin filament formation (*CEP250* KO), rootletin-CEP68 filaments (*CROCC* KO, *CEP68* KO), and microtubules attached to subdistal appendages (*CEP128* KO). We also analyzed *CEP128-250* and *CEP128-68* double KO cells to impair all three centrosome cohesion pathways or the subdistal appendage derived microtubules and the rootletin-CEP68 filaments, respectively. *NIN* KO cells were excluded from this analysis because of the multifunctional roles of ninein in microtubule anchorage at centrosomes, centrosome cohesion, and the recruitment of rootletin–CEP68 complexes to centrosomes. Furthermore, subdistal appendages remain present in *NIN* KO cells, as observed by EM (Fig. EV1A). *AKAP450* KO cells were also excluded because AKAP450 has additional Golgi-associated functions that are independent of centrosome cohesion mechanisms.

In 80% of wild-type RPE1 cells, both centrosomes overlapped with the Golgi marker GM130 (Fig EV3A and B), which is consistent with the STED analysis (Appendix Fig. 2C). Deletion of *CEP250*, *CROCC*, *CEP68* or *CEP128* increased the proportion of cells in which at least one centrosome was no longer associated with the Golgi, from approximately 20% in wild-type cells to 40–60% (Fig EV3B). This suggests that probably all three elements of centrosome cohesion are important for the attachment of centrosomes to the Golgi. The combined deletion of *CEP128* and *CEP250* (*CEP128-250*) had the strongest effect, with approximately 80% of cells exhibiting at least one centrosome that was no longer associated with the Golgi (Fig EV3A and B). *CEP128*-*250* KO cells lack all elements of centrosome cohesion and therefore it is not surprising that the impact on centrosome-Golgi co-localization is the strongest. *CEP128-CEP68* KO cells behaved similarly to *CEP68* KO and *CEP128* KO cells, suggesting that rootletin–CEP68 filaments and subdistal appendage microtubules function in a linear pathway to mediate centrosome-Golgi anchoring.

To validate the findings from the fixed-cell analysis and monitor changes in centrosome-Golgi distance over time, we measured centrosome-Golgi association in living RPE1 cells expressing γ-tubulin–mRuby2 as a centrosome marker and GalT–GFP as a Golgi marker (Cole *et al*., 1996). We tracked 15 individual cells over an 80-minute time course. In wild-type RPE1 cells, both centrosomes co-localized with the Golgi marker in 67% of cells throughout the 80-min live-cell imaging period (Fig 5A and B, yellow bars). In two cells, both centrosomes did not overlap with the GalT-GFP marker over the 80 min analysis (Fig 5A and B, purple bars). However, even in cases of detachment, centrosomes still localized close to the Golgi with a maximum distance of 2.3 µm (Fig 5B, Movie EV3).

**Figure 5.**
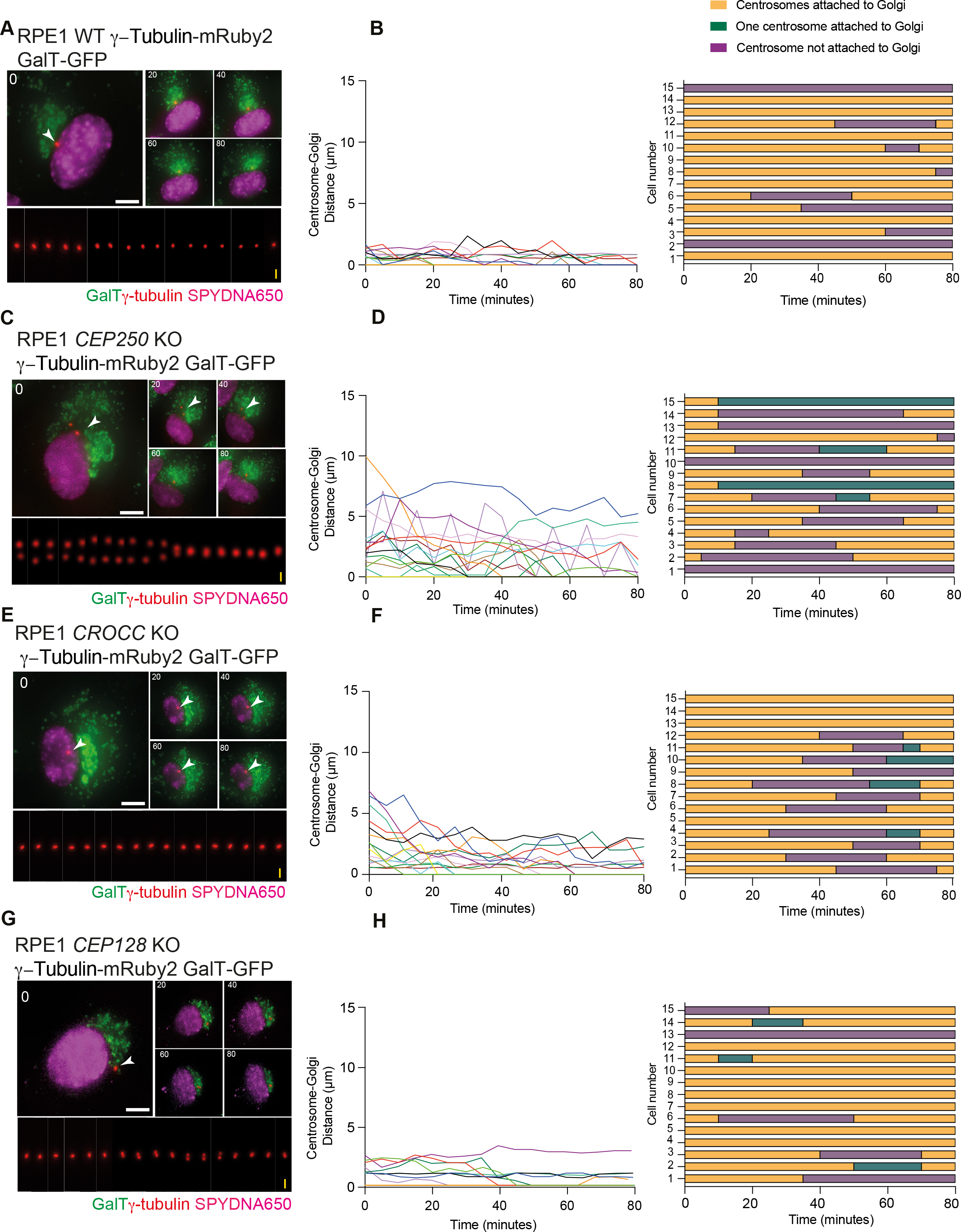
Spatial distribution of the centrosome and the Golgi in WT and linker mutants. **A, C, E, G** Time-lapse images (5 min interval) of RPE1 WT, *CEP250* KO, *CROCC* KO, *CEP128* KO cells expressing γ-tubulin-mRuby2 (red) as markers for the centrosomes and GalT-GFP as a marker for the Golgi. Representative still images (top rows) and kymographs of centrosomes (bottom rows) are shown. The DNA was marked with SPY DNA650 (magenta). White scale bars, 5 µm. Yellow scale bars (vertical), 2 µm. (Movie EV3). **B, D, F, H** Centrosome-Golgi distances (µm) from (A), (C), (E) and (G) are plotted over time. For cells with separated centrosomes, the mean distance from each centrosome to the Golgi was measured Timeline for individual cells (n=15 cells, pooled from N=3 independent experiments) monitored over 80 minutes. Orange bars indicate when the centrosome is attached to the Golgi; purple bars indicate the lack of attachment of the centrosome to Golgi; green bars indicate when one centriole is attached to the Golgi. Each row corresponds to one cell, and color transitions mark the time of attachment change.

*CEP250* KO cells, which are defective in both centrosome cohesion and the centrosomal attachment of rootletin–CEP68 filaments, displayed the strongest defect among the mutants in centrosome-Golgi attachment. During the 80-min live-cell imaging period, all 15 centrosomes lost co-localisation with the Golgi and subsequently partially re-associated with it. In most cases both centrosomes detached from the Golgi; in less frequent cases, only one centrosome lost co-localisation with the GalT-GFP signal (Fig. 5C and D; green bars). On average, centrosomes were 2 µm away from the Golgi. However, peak values were 7-10 µm (Fig 5C and D, Movie EV3).

*CROCC* KO cells exhibited an intermediate phenotype compared with wild-type and *CEP250* KO cells. Similar to *CEP250* KO cells, the most prominent phenotype in *CROCC* KO cells was the detachment of both centrosomes. However, detachment periods were shorter, and, in cases of detachment, peak values were reduced by 6 µm compared with *CEP250* KO cells (Fig 5E and F, Movie EV3). The phenotype of *CEP128* KO cells was the least severe of the three KO cells. These cells behaved similarly to WT cells; however, when the centrosomes detached from the Golgi, the distance between the centrosomes and the Golgi was greater than that observed in WT cells (Fig. 5G and H, Movie EV3).

Taken together, these findings demonstrate that all three centrosome cohesion mechanisms contribute to centrosome-Golgi association. The subdistal-derived microtubules had the least pronounced effect. The data suggest that centrosome-Golgi attachment is a largely stable state punctuated by brief, periods of partial or full disengagement across the cell population.

### The centrosome linker facilitates Golgi reformation

We investigated how centrosome cohesion elements influence the spatial arrangement of the Golgi apparatus. In RPE1 cells, the Golgi is positioned near the nuclear envelope and often overlaps with the centrosome. Previous work has reported that the Golgi becomes more stretched along the nuclear envelope as cells transition from G1 to S phase (Frye *et al*., 2020). Using EdU (S phase labelling) and the centrosome distance as markers, we observed compact and extended Golgi at the nuclear envelope in G1 phase. The compact Golgi was more pronounced in S phase. In late G2 (separated centrosomes) the Golgi was mainly extended along the nuclear envelope (Fig EV4A and B). In *CEP250* KO cells, the compact and extended Golgi states were similarly distributed across G1 and S phase (Fig. EV4A and B) indicating that loss of *CEP250* influences Golgi morphology.

To determine how the different centrosome cohesion pathways affect Golgi morphology, we compared the Golgi in G1/S-phase in wild-type and centrosome cohesion mutant RPE1 cells. Cells were fixed and stained for the Golgi marker GM130 and γ-tubulin to visualize centrosomes. CENPF staining was additionally used to distinguish G1/S-phase cells from G2 cells. The distribution of GM130 relative to the DAPI-stained nucleus was classified into three categories: compact, half-moon, and compact morphology (Fig EV4C). In wild-type RPE1 cells, the Golgi predominantly displayed a compact morphology as observed in Fig EV4B. *CEP128* KO cells showed a similar distribution of Golgi morphologies to wild-type cells (Fig EV4C), whereas *CEP250* KO, *CROCC* KO, and *CEP68* KO cells exhibited an increased frequency of the extended Golgi configuration (Fig EV4C). In all conditions, the Golgi remained organized around the nuclear envelope, indicating that this association is not affected by loss of centrosome cohesion components (Fig EV4C). This suggests that rootletin–CEP68 filaments, but not microtubules at the subdistal appendages, influence Golgi morphology.

We next investigated the role of centrosome cohesion elements in Golgi reformation. We disrupted Golgi structure by treating wild-type and mutant cells for 2 hours with the drug brefeldin A, a reversible inhibitor of vesicular transport between the endoplasmic reticulum and the Golgi (Misumi *et al*., 1986). DMSO, the brefeldin solvent, was used as control. After washing out brefeldin, we monitored Golgi reformation after 30 min and 2 hours using GM130 staining as Golgi marker (Fig 6A and B). Golgi reformation was quantified by measuring the radial distribution of GM130 signal intensity from the Golgi center (Fig 6C). In wild-type cells, centered Golgi distribution was visibly restored within 30 min after drug wash out (Fig 6B and C). In contrast, recovery was slower in *CEP250*, *CROCC*, and *CEP68* cells (Fig 6B and C). *CEP128* KO cells showed slightly faster recovery after 30 min compared with the other mutants, but still remained slower than in WT cells. Notably, *CEP128-250* and *CEP128-68* double KO cells (Fig EV4D and E) behaved similarly to the respective *CEP250* and *CEP68* single KO. The Golgi assembly defect of *CEP250* KO cells was rescued by expression of *CEP250* (Fig EV4D and E).

**Figure 6.**
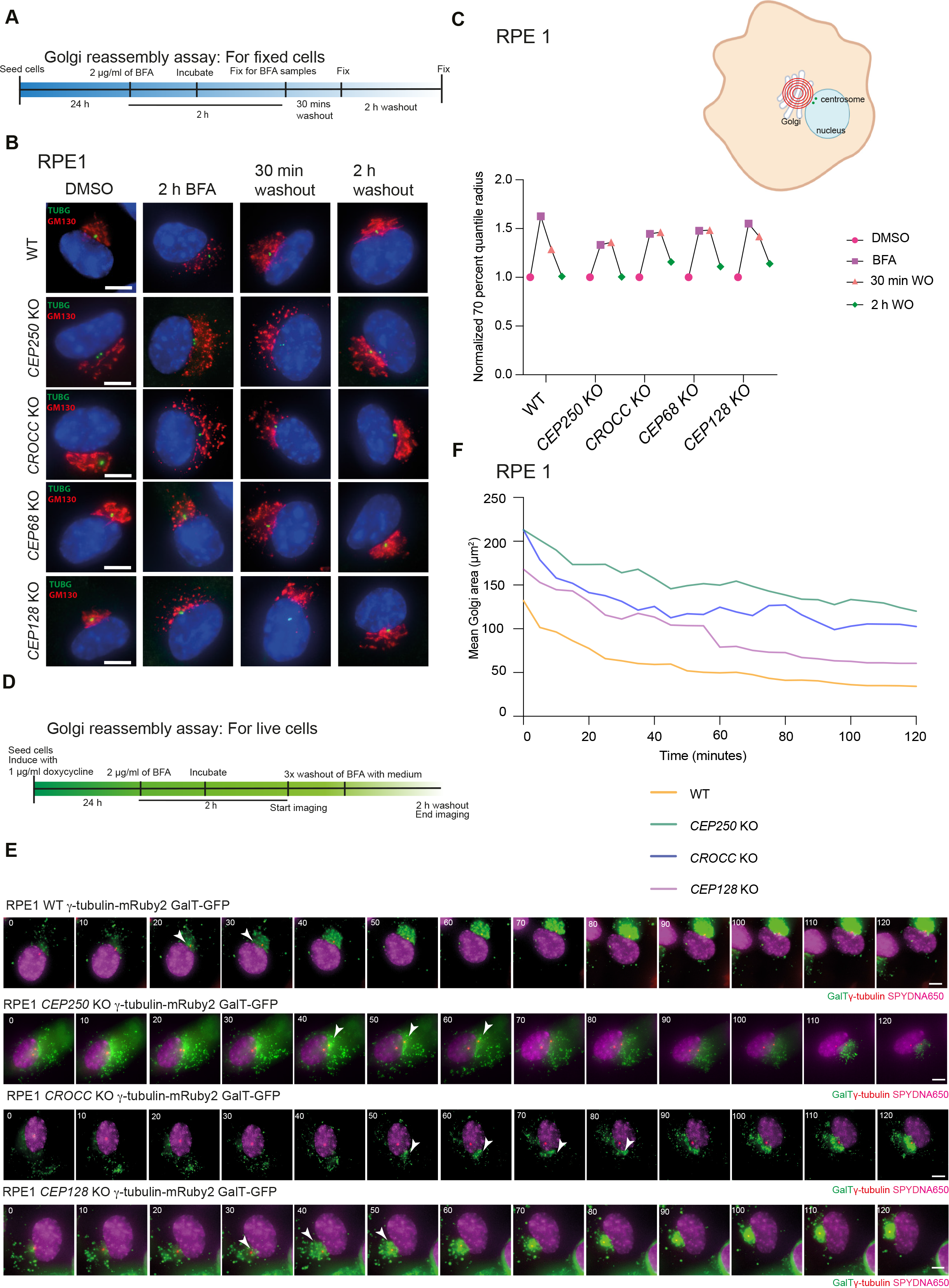
Reformation of Golgi stacks after brefeldin washout is dependent on the presence of linker proteins. **A** Experiment design for Golgi reassembly assay for fixed cells. Cells were treated with 2 µg/ml of BFA for 2 h. Cells were then fixed at three different time points: two hours after drug treatment, 30 minutes after washout and 120 minutes after washout. **B** Representative fluorescent images of RPE 1 WT, *CEP250* KO, *CROCC* KO, *CEP68* KO and *CEP128* KO cells. γ-tubulin (green) and GM130 (red) antibodies were used for indirect fluorescent analysis. The cells were imaged at three different timepoints; two hours after drug treatment, 30 minutes after washout and 120 minutes after washout. DMSO treated cells were used as control. DNA was stained with DAPI. Scale bars: 5 µm. **C** Analysis of Golgi distribution from cells in (C). Radius of a circle that was drawn around the centre of mass of the cell’s Golgi signal (GM130 signal) such that the circle encloses 70% of the total signal intensity of the GM130 signal. N=3 independent experiments; n≥80 cells. Faster accumulation (smaller diameter) means more dense distribution. **D** Experiment design for Golgi reassembly assay for live cells. Cells were treated with 2 µg/ml of BFA for 2 h. Drug was washed out and reformation of Golgi was observed. **E** Time-lapse images (5 min interval) of RPE1 WT, *CEP250* KO and *CROCC* KO, *CEP128* KO cells expressing γ-tubulin-mRuby2 (red) as markers for the centrosomes with and GalTGFP as marker for the Golgi. Shown are representative still images (10 min interval). White arrows indicate formation of first Golgi stack. The DNA was marked with SPY DNA650 (magenta). White scale bars, 5 µm. (Movie EV4). **F** Area of the Golgi was measured over 120 minutes (µm^2^) from (E) plotted over time (n = 15 cells, pooled from N = 3 independent experiments).

We also analyzed the co-localization of centrosomes with the Golgi 2 h after brefeldin A washout. In >80% of wild-type cells, both centrosomes co-localized with the Golgi area (Fig. EV4F). In *CEP250* KO, *CROCC* KO, and *CEP68* KO cells, approximately 40–50% of cells showed no co-localization of both centrosomes with the Golgi. In contrast, *CEP128* KO cells behaved similarly to wild-type cells (Fig. EV4F). These results suggest that, in this Golgi reformation assay, microtubules associated with subdistal appendages do not contribute to the reestablishment of centrosome-Golgi attachment.

The findings from the fixed-cell analysis of centrosome reformation were confirmed by live-cell imaging of RPE1 cells expressing γ-tubulin-mRuby2 and GalT-GFP (Fig 6D-F). In this experiment, we quantified Golgi area compaction over time (Fig 6F, Movie EV4). Before brefeldin A washout, the Golgi area was more compact in RPE1 wild-type and *CEP128* KO cells than in *CEP250* KO and *CROCC* KO cells (Fig 6E and F). After drug wash out, the compaction over time was strongest in wild type and *CEP128* KO cells, while recovery was slower for *CROCC* and *CEP250* KO cells (Fig 6F, Movie EV4).

Together, these results demonstrate that all three centrosome cohesion mechanisms contribute to Golgi compaction and reformation. Centrosome distance and rootletin filaments have the strongest impact on Golgi reformation, while subdistal appendage-derived microtubules have the least effect.

### Centrosome cohesion mechanisms affect microtubule re-formation and microtubule acetylation

Golgi reassembly following microtubule depolymerization and repolymerization is accompanied by the rapid formation of Golgi-associated acetylated microtubules, which are an indication for stable microtubules (Thyberg & Moskalewski,1993). We therefore investigated whether microtubule density and tubulin acetylation at the Golgi were altered 5 min after microtubule reassembly, following cold-induced depolymerization and subsequent rewarming, in wild-type and linker-mutant cells (Fig. EV5A). Microtubule density at the Golgi, marked by GM130, was reduced in *CEP250* KO, *CROCC* KO, *CEP68* KO, *CEP128* KO, *CEP128-250* KO, and *CEP128-68* KO cells compared with wild-type cells (Appendix Fig. S8A and B). In addition, acetylated microtubule formation at both the centrosome and Golgi was also markedly decreased in these mutant cells 5 min after microtubule reformation (Fig. EV5B and C). These findings indicate that all centrosome cohesion elements contribute to microtubule reformation, Golgi-associated microtubule organization, and microtubule acetylation (Thyberg & Moskalewski, 1993).

We also examined centrosome positioning after 5 min of microtubule regrowth. Upon microtubule reformation, both centrosomes overlapped with the Golgi in 85% of wild-type cells. This proportion decreased to approximately 70% in *CEP250* and *CEP128* KO cells, whereas only minor effects were observed in *CROCC* and *CEP68* KO cells (Fig EV5D). These findings suggest that centrosome proximity and subdistal appendage-derived microtubules contribute to centrosome repositioning toward the Golgi following microtubule repolymerization. Notably, this mechanism appears to differ from Golgi reformation following brefeldin A washout, during which subdistal appendages play a more limited role (Fig. 6).

### Centrosome cohesion elements affect cell migration in distinct ways

The centrosome-Golgi axis is well established as an important regulator of cell migration (Wakida *et al*., 2010). However, how the different mechanisms of centrosome cohesion affect cell migration remains unknown. To address this, we performed wound-healing assays of RPE1 cells defective in centrosome cohesion mechanisms (Fig 7A and B, Fig EV5E and F). In this assay, cell migration was most severely impaired in *CEP250* and *CEP128* KO cells followed by *CEP68* and *CROCC* KO cells. *AKAP450* KO cells, which were included as a control, showed a relatively mild defect in the wound-healing assay, whereas the cell migration defect observed in *CEP250* KO cells was rescued by *CEP250* expression (Fig EV5E and F). In addition, *CEP128-68* and *CEP128-250* double KO cells exhibited a more severe migration defect than the single mutants (Fig 7A and B), indicating that the different elements of centrosome cohesion function in parallel rather than in a linear pathway. We also tracked the migration of individual cells toward the closing wound, with a particular focus on migration speed and directional persistence (Fig 7C-E). Mean cell velocity was reduced in *CEP250* KO, *CROCC* KO, *CEP68* KO, and *CEP128* KO cells, with the strongest reduction among the single KOs observed in *CEP128* KO cells (Fig 7C and D). *CEP128-250* KO cells exhibited the strongest reduction in mean velocity, whereas *CEP128-68* KO cells behaved similarly to *CEP68* KO cells but showed a less severe phenotype than *CEP128* KO cells, indicating partial rescue of the *CEP128* KO defect. This rescue may be attributed to an as-yet-unclear function of CEP68, which, unlike rootletin, also localizes to the Golgi (Hung *et al*., 2026).

**Figure 7.**
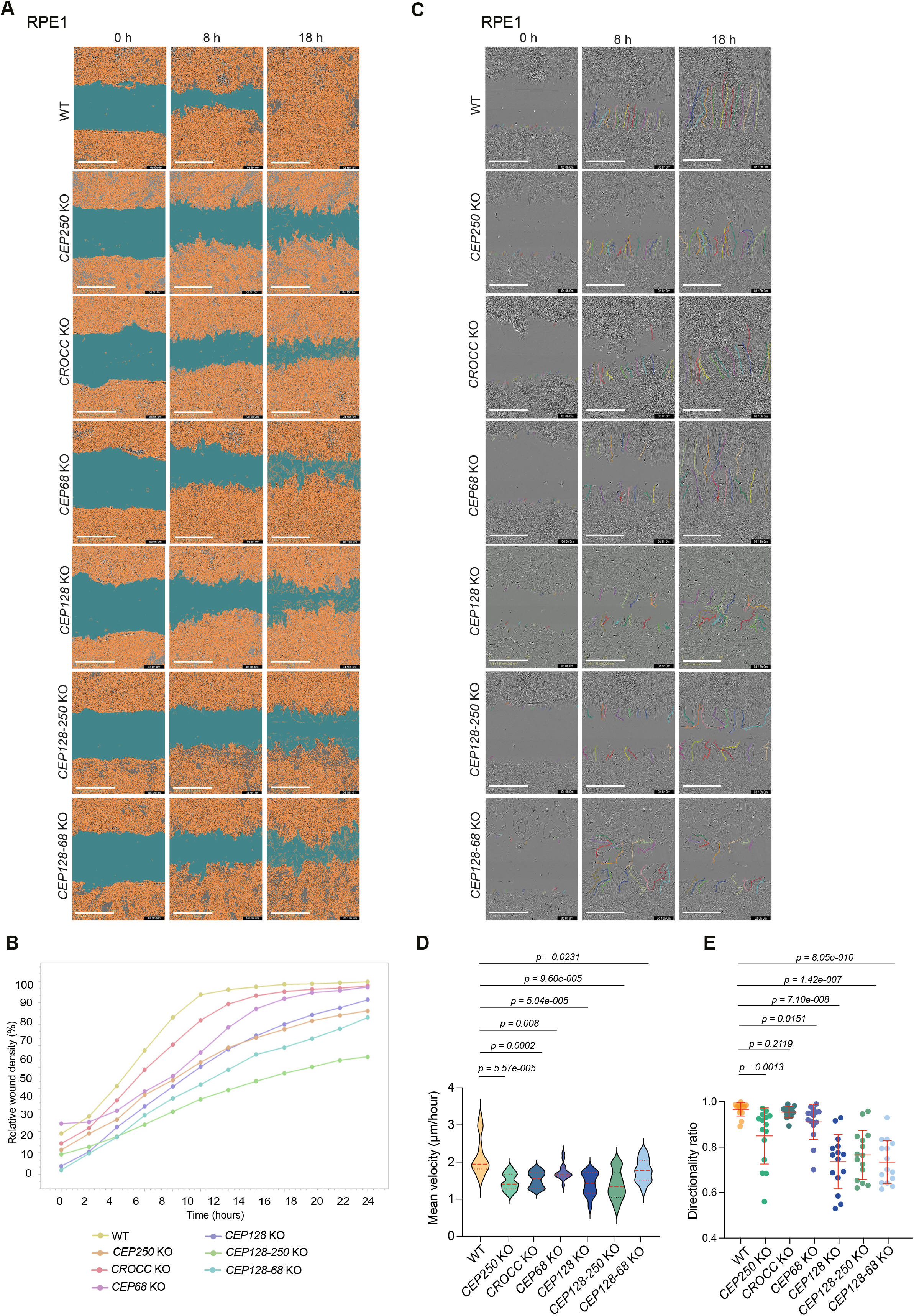
Effect of centrosome linker mutants on scratch wound closure. **A** Representative image of scratch wound analysis of WT, *CEP250* KO, *CROCC* KO, *CEP68* KO, *CEP128* KO, *CEP128-250* KO and *CEP128-68* KO cells. Phase contrast overlaid images illustrate wound gaps at 0 h, 8 h and 18 h. Scale bars: 600 µm. **B** Relative wound closure of the indicated cell lines over 24 h, measured using the Incucyte Scratch Wound assay. Wound closure is expressed as relative wound density (%), normalized to the initial wound at 0 h. Images were acquired every 2 h with a 10× objective. n=15 cells pooled from N=3 independent experiments. **C** Representative overlayed image of single cell tracking of WT, *CEP250* KO, *CROCC* KO, *CEP68* KO, *CEP128* KO, *CEP128-250* KO and *CEP128-68* KO cells. Phase contrast images illustrate wound gaps at 0 h, 8 h and 18 h and colored lines represent the movement of the single cells. Scale bars: 600 µm. **D** Mean velocity of single cells tracked during wound-closure migration. Cells at the wound edge (from (C)) were tracked over 24 h (images every 2 h, 10× objective) using MTrackMateJ in Fiji. Mean velocity (µm/hour) was calculated per cell as total path length divided by tracking time. Violin plots show the kernel density distribution of all tracked cells; the embedded box plots show the median (center line), interquartile range (box), and 1.5× IQR (whiskers). Individual points represent single cells n=15 cells, pooled from N=3 independent experiments. Unpaired t test. **E** Directionality ratio of single cells (same cells as in (D)) during wound-closure migration. Cells at the wound edge were tracked over 24 h (images every 2 h, 10× objective) using MTrackMateJ in Fiji. The directionality ratio was calculated for each cell as the distance between the start and end points of the track divided by the total path length, where 1 indicates a perfectly straight path and values approaching 0 indicate random movement. n=15 cells pooled from N=3 independent experiments. Unpaired t test.

Analysis of migration directionality revealed that *CEP128* KO cells also exhibited the most severe defect, followed by *CEP250* KO, *CEP68* KO, and *CROCC* KO cells (Fig 7E). *CEP128-250* KO and *CEP128-68* KO cells displayed directionality defects similar to those observed in *CEP128* KO cells, with *CEP128-68* KO cells showing a slightly more pronounced phenotype. The combination of slightly higher mean velocity and reduced directionality in *CEP128-68* KO cells may explain the reduced relative wound density observed in these cells compared to the single KO cell lines (Fig 7B, D, and E). These findings suggest that different components of the centrosome cohesion machinery regulate distinct aspects of cell migration. Loss of *CEP128* had the strongest effect on both migration speed and directional persistence. Disruption of the rootletin–CEP68 filament network reduced migration speed but did not impair directionality.

## Discussion

Rootletin–CEP68 filaments were originally described as components of ciliary rootlets (Yang *et al*., 2002). Using U2OS cells and HCT116 cells, it was then found that rootletin forms filaments together with the bundling protein CEP68 at the centrosome that function in centrosome cohesion (Bahe *et al*., 2005; Vlijm *et al*., 2018; Graser *et al*., 2007). However, in RPE1 cells rootletin-CEP68 filaments do not contribute to centrosome cohesion despite remaining attached to both the mother and daughter centrioles and forming the same periodic structure with CEP68 as in HCT116 cells (Theile *et al*., 2023 ; Vlijm *et al*., 2018). These observations raise the question of whether the rootletin-CEP68 filaments have a function beyond mediating centrosome cohesion.

Here, we report that rootletin-CEP68 filaments at the centrosome function in Golgi organization. Rootletin-CEP68 filaments recruit AKAP450 through CEP68 and extend into the Golgi region where AKAP450 interacts with GM130 (Rivero *et al*., 2009). Consistent with this model, detached rootletin filaments align with Golgi filaments in 99% of *CEP250* KO cells containing rootletin filaments. In addition, cells lacking rootletin show a reduced frequency of centrosome-Golgi association compared with control cells. Furthermore, the absence of either rootletin or CEP68 delays Golgi reassembly following brefeldin A washout and impairs cell migration, which depends on proper Golgi organization.

Previously, AKAP450 was mainly described as a PCM associated protein. This conclusion is largely based on the PCM localization of the overexpressed C-terminal PACT domain construct (Gillingham & Munro, 2000). Also, in this analysis, the isolated AKAP450^PACT^ domain predominantly localized to γ-tubulin-marked centrioles where it probably interacts with CEP57, a known interactor of the PACT domain of PCNT (Watanabe *et al*., 2020). Surprisingly, AKAP450^PACT^ showed only a sparse association with rootletin-CEP68 filaments, in contrast to full-length AKAP450, which predominantly localized to these filaments rather than to the γ-tubulin-marked PCM. Furthermore, comparison of AKAP450 centrosome localization between interphase and metaphase centrosomes revealed a strong decline of AKAP450 at mitotic centrosomes, which lack the rootletin filaments (Fry *et al*., 1998), further supporting the localization of AKAP450 predominantly with rootletin filaments. These findings therefore suggest that regions of AKAP450 beyond the PACT domain are required for efficient targeting of the full-length protein to rootletin–CEP68 filaments probably by suppressing its association with the PCM. Thus, at least in the cell lines examined, AKAP450 is not a major PCM component involved in centrosomal microtubule nucleation through CDK5RAP2 recruitment. This function is likely mediated by the Golgi-associated pool of AKAP450, which also carries CDK5RAP2 (Gavilan *et al*., 2018; Wang *et al*., 2010).

The role of rootletin–CEP68–AKAP450 filaments in Golgi organization is embedded within a broader network of proteins that collectively promote centrosome cohesion (Hata *et al*., 2019; Theile *et al*., 2023). Firstly, the rootletin–CEP68–AKAP450 filaments directly connect the centrosome to the Golgi. Since AKAP450 associates with the rootletin–CEP68 filaments even when they are not attached to the Golgi (Appendix Fig. S2C), AKAP450-bound rootletin filaments may assemble first and then interact with GM130 at the Golgi. Alternatively, rootletin–CEP68 may interact with AKAP450 that is already associated with GM130. Secondly, the C-Nap1–ninein linker regulates the distance between the two centrosomes, anchors rootletin-CEP68 filaments to centrosomes and also has a not well-defined role in microtubule nucleation (Theile *et al*., 2023). Thirdly, microtubules originating from the subdistal appendages also maintaining the close proximity between the two centrosomes (Hata *et al*., 2019). In addition, these microtubules may contribute to Golgi organization either through direct interactions with the Golgi apparatus or by interacting with Golgi-derived microtubules, potentially via the kinesin motor KIFC3, which associates with subdistal appendage-derived microtubules and functions in Golgi positioning (Xu *et al*., 2002;Hata *et al*., 2019). The observation that the centrosome-Golgi attachment defect in *CEP250* knockout cells is exacerbated by removal of the subdistal appendage microtubules (Fig. 5) (Kashihara *et al*., 2019) further supports the role of subdistal appendage attached microtubules in Golgi organization. Together, these findings revise the current model of how the centrosome organizes the Golgi. Rather than relying only on PCM/centrosome-derived microtubules (Mascanzoni *et al*., 2022), Golgi organization emerges from the coordinated action of the microtubule cytoskeleton, including the specialized microtubules of the subdistal appendages, and the rootletin-CEP68-AKAP450 filament network and the centrosome distance that likely regulates the local density of the rootletin-CEP68 fibres and thereby the extent of centrosome-Golgi connectivity.

Our analysis focuses on the centrosome-Golgi connection by cohesion mechanisms in dividing cells. However, centrosome cohesion has also been implicated in serum starvation-induced ciliogenesis, during which subdistal appendages contribute to the spatial assembly of the primary cilium either in the cytoplasm or exposed on the cytoplasmic membrane (Mazo *et al*., 2016). We propose that perturbation of the mechanisms described in this manuscript that regulate centrosome-Golgi attachment in cycling cells may determine the intracellular site of primary cilium assembly during ciliogenesis.

The effects of the three centrosome cohesion mechanisms on cell migration are particularly interesting. During cell migration, Golgi dynamics are tightly regulated, with the Golgi apparatus repositioning toward the leading edge of the cell to support polarized trafficking and secretion. This process involves the fragmentation, redistribution, and subsequent reassembly of the Golgi in the direction of migration. Notably, key aspects of Golgi remodeling during migration, including Golgi positioning, compaction, and reformation, are differentially controlled by the three centrosome cohesion mechanisms already suggesting distinct phenotypes (Table 6). Previous work showed that disruption of AKAP450 through overexpression of its N-terminal GM130-binding domain impairs both Golgi organization and cell migration (Hurtado *et al*., 2011). Because AKAP450 has multiple functions at both the centrosome and the Golgi, the precise basis of the migration phenotype has remained unclear. However, deletion of the rootletin-CEP68 fibers has only a moderate effect on cell migration compared with wild-type cells, as assessed by wound closure as well as the mean velocity and directionality of individual cells (Fig 7F). Therefore, it is unlikely that the strong migration defect observed in AKAP450-displaced cells is caused by the loss of rootletin-CEP68 fiber attachment to the Golgi. Rather it suggests that microtubules anchored to the Golgi via AKAP450 have an important function in cell migration.

**Table 1.**
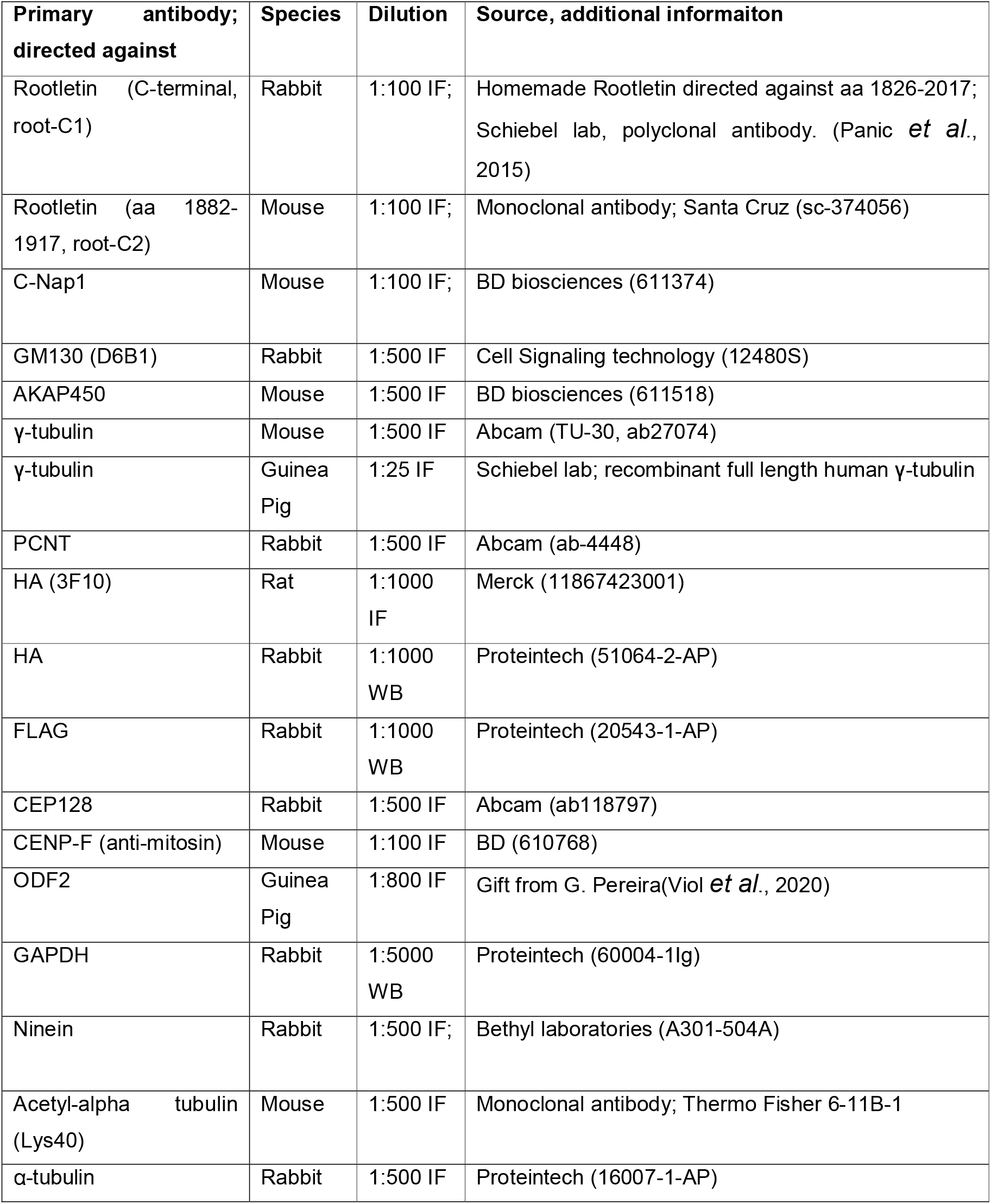
Primary antibodies.

| Primary antibody;<br>directed against | Species | Dilution | Source, additional informaiton |
| --- | --- | --- | --- |
| Rootletin (C-terminal, root-C1) | Rabbit | 1:100 IF; | Homemade Rootletin directed against aa 1826-2017; Schiebel lab, polyclonal antibody. (Panic *et al.*, 2015) |
| Rootletin (aa 1882-1917, root-C2) | Mouse | 1:100 IF; | Monoclonal antibody; Santa Cruz (sc-374056) |
| C-Nap1 | Mouse | 1:100 IF; | BD biosciences (611374) |
| GM130 (D6B1) | Rabbit | 1:500 IF | Cell Signaling technology (12480S) |
| AKAP450 | Mouse | 1:500 IF | BD biosciences (611518) |
| γ-tubulin | Mouse | 1:500 IF | Abcam (TU-30, ab27074) |
| γ-tubulin | Guinea Pig | 1:25 IF | Schiebel lab; recombinant full length human γ-tubulin |
| PCNT | Rabbit | 1:500 IF | Abcam (ab-4448) |
| HA (3F10) | Rat | 1:1000 IF | Merck (11867423001) |
| HA | Rabbit | 1:1000 WB | Proteintech (51064-2-AP) |
| FLAG | Rabbit | 1:1000 WB | Proteintech (20543-1-AP) |
| CEP128 | Rabbit | 1:500 IF | Abcam (ab118797) |
| CENP-F (anti-mitosin) | Mouse | 1:100 IF | BD (610768) |
| ODF2 | Guinea Pig | 1:800 IF | Gift from G. Pereira(Viol *et al.*, 2020) |
| GAPDH | Rabbit | 1:5000 WB | Proteintech (60004-1Ig) |
| Ninein | Rabbit | 1:500 IF; | Bethyl laboratories (A301-504A) |
| Acetyl-alpha tubulin (Lys40) | Mouse | 1:500 IF | Monoclonal antibody; Thermo Fisher 6-11B-1 |

| Primary antibody;<br>directed against | Species | Dilution | Source, additional information |
| --- | --- | --- | --- |
| α-tubulin | Rabbit | 1:500 IF | Proteintech (16007-1-AP) |

**Table 2.** Secondary antibodies.

| Secondary antibody | Species | Dilution | Source |
| --- | --- | --- | --- |
| Anti-mouse IgG Alexa Fluor 488/555/647 | Goat | 1:500 IF | Thermo Fisher scientific;<br>add number in all cases |
| Anti-rabbit IgG Alexa Fluor 488/555/647 | Goat | 1:500 IF | Thermo Fisher scientific |
| Anti-goat IgG Alexa Fluor 488/555/647 | Goat | 1:500 IF | Thermo Fisher scientific |
| Anti-guinea pig IgG Alexa Fluor 555/647 | Goat | 1:500 IF | Thermo Fisher scientific |
| Anti-guinea pig IgG Alexa Fluor 488 | Donkey | 1:500 IF | Sigma-Aldrich |
| Anti-mouse IgG STAR /580/635P | Goat | 1:100 IF | Abberior |
| Anti-rabbit IgG STAR 580/635P | Goat | 1:100 IF | Abberior |
| Anti-rat STAR RED | Goat | 1:100 IF | Abberior |
| Anti-rat IgG STAR 580/635P | Goat | 1:100 IF | Abberior |
| Anti-mouse HRP | Goat | 1:5000 WB | Jackson ImmunoResearch |
| Anti-rabbit HRP | Donkey | 1:5000 WB | Jackson ImmunoResearch |
| Anti-rat HRP | Goat | 1:5000 WB | Thermo Fisher scientific |

**Table 3.**
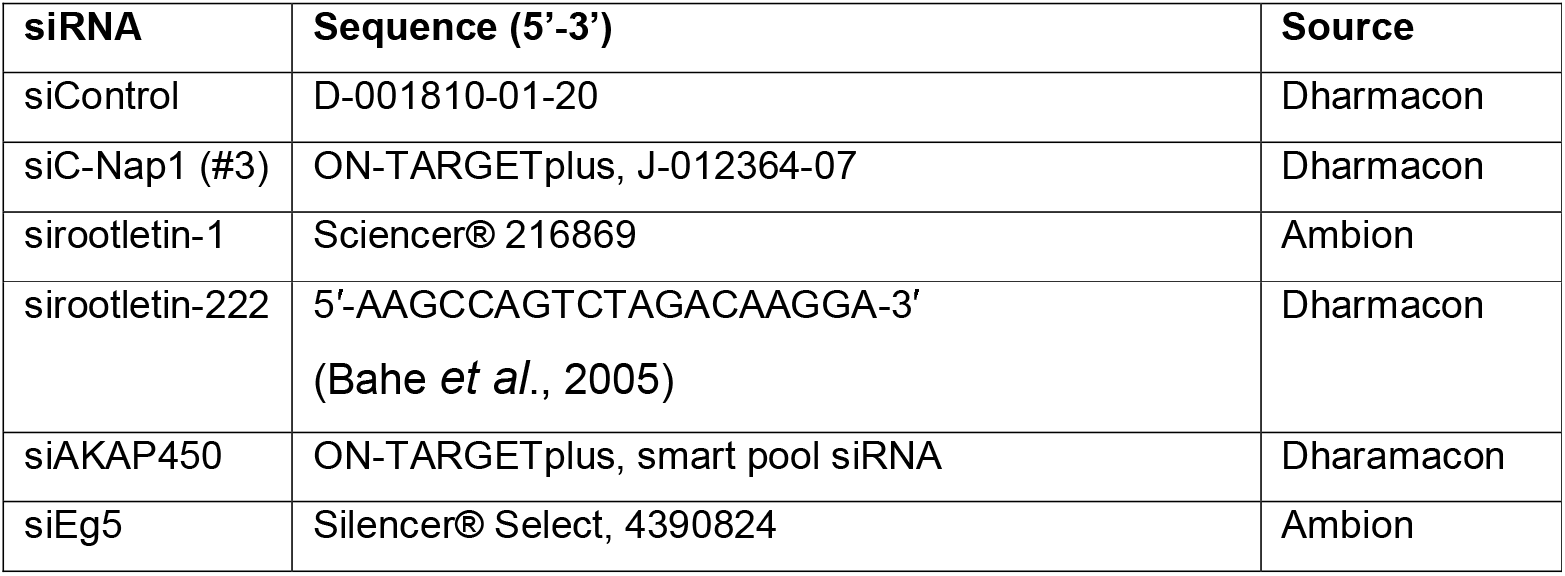
siRNAs.

**Table 4.**
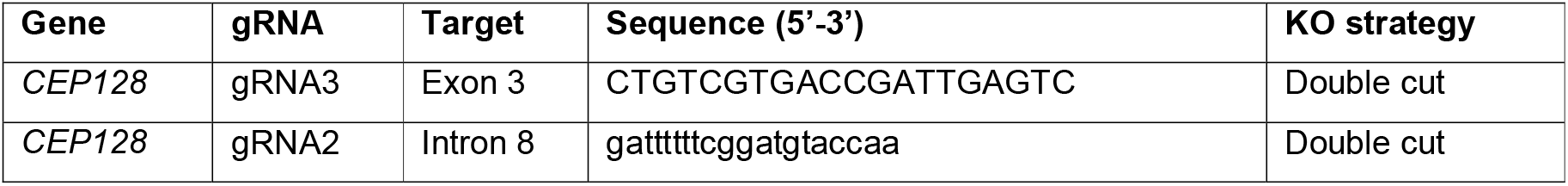
gRNAs for knock-out generation.

**Table 5.** Cell lines.

| Cell line | Source/reference |
| --- | --- |
| RPE1 tetON WT | Schiebel lab |
| RPE1 tetON *CEP250* KO #17 | (Panic *et al.*, 2015) |
| RPE1 tetON *CEP128* KO #20 | Generated in this study |
| RPE1 tetON *CEP128* -*CEP250* KO #57 | Generated in this study |
| RPE1 tetON *CEP128* -*CEP68* KO #109 | Generated in this study |
| RPE1 tetON *NIN* KO #14 | (Theile *et al.*, 2023) |
| RPE1 tetON *CROCC* KO #49 | (Theile *et al.*, 2023) |
| RPE1 tetON *CEP68* KO #20 | (Panic *et al.*, 2015) |
| RPE1 *AKAP450* KO | (Gavilan *et al.*, 2018) |
| HCT116 tetON WT | (Theile *et al.*, 2023) |
| HCT116 tetON *CEP250* KO #234 | (Theile *et al.*, 2023) |
| HCT116 tetON *CROCC* KO #79 | (Theile *et al.*, 2023) |
| HCT116 tetON *NIN* KO #139 | (Theile *et al.*, 2023) |
| RPE1 tetON WT PACT<sup>AKAP</sup>-HA | Generated in this study |
| RPE1 tetON *CEP250* KO #17 PACT<sup>AKAP</sup>-HA | Generated in this study |
| RPE1 tetON *CEP68* KO #20 PACT<sup>AKAP</sup>-HA | Generated in this study |
| RPE1 tetON WT PACT<sup>PCNT</sup>-HA | Generated in this study |
| RPE1 tetON *CEP250* KO #17 PACT<sup>PCNT</sup>-HA | Generated in this study |
| RPE1 tetON *CEP68* KO #20 PACT<sup>PCNT</sup>-HA | Generated in this study |
| RPE1 tetON WT γ-tubulin-mRuby2 GalT-GFP | Generated in this study |
| RPE1 tetON *CEP250* KO #234 γ-tubulin-mRuby2 GalT-GFP | Generated in this study |
| RPE1 tetON *CROCC* KO #79 γ-tubulin-mRuby2 GalT-GFP | Generated in this study |
| RPE1 tetON *CEP128* KO #20 γ-tubulin-mRuby2 | Generated in this study |
| RPE1 tetON *CEP128* KO #20 γ-tubulin-mRuby2 GalT-GFP | Generated in this study |
| HCT 116 tetON WT γ-tubulin-mRuby2 | Generated in this study |
| HEK293 T | Schiebel lab |
| HEK293 GP | Schiebel lab |
| U2OS tetON | Schiebel lab |

**Table 6.**
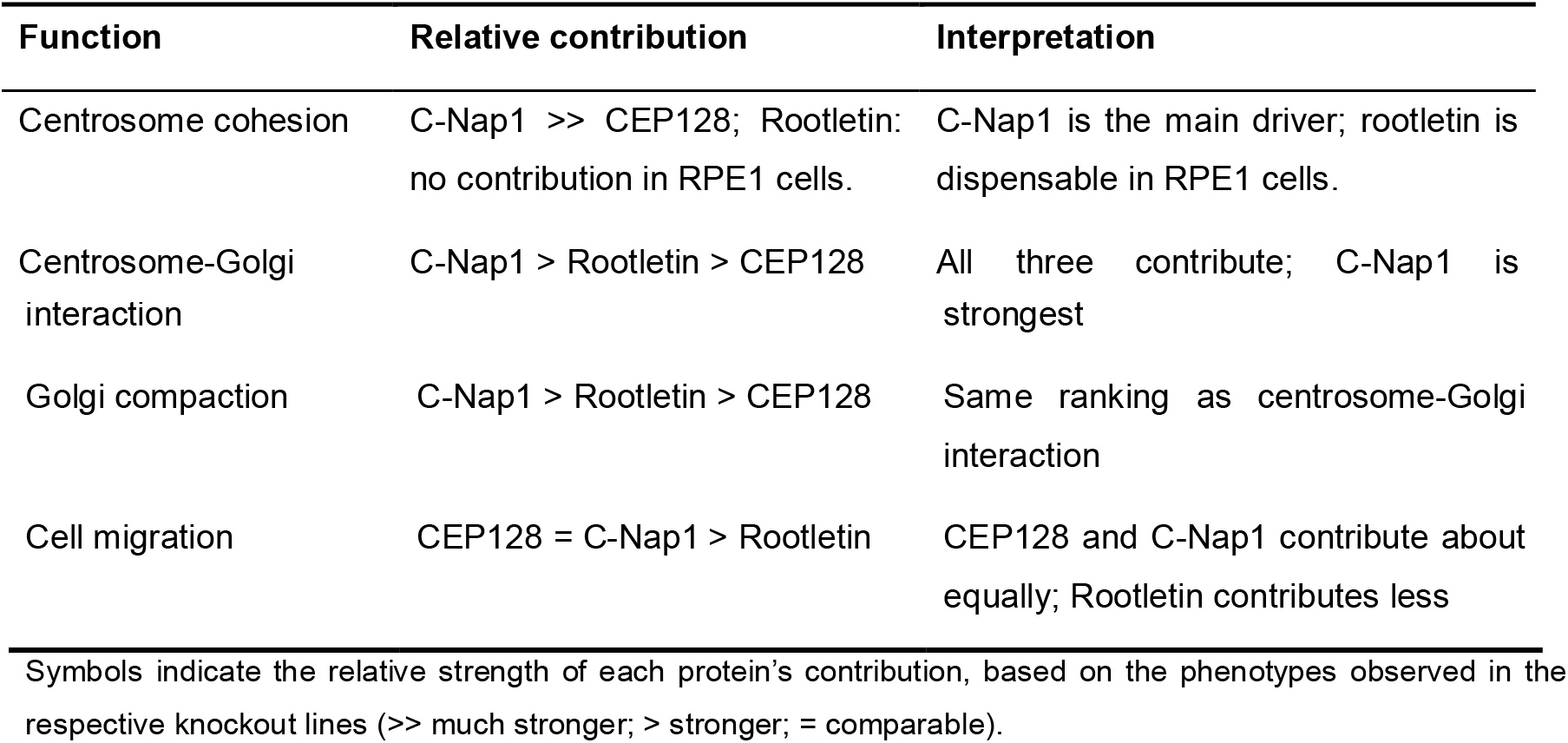
Relative contributions of C-Nap1, CEP128 and Rootletin (CROCC) to centrosome-associated functions.

In contrast to *CROCC* KO and *CEP68* KO cells, loss of *CEP250* or *CEP128* resulted in a more pronounced impairment of cell migration parameters, suggesting that disruption of the rootletin-CEP68 connection per se is less important for cell migration than the defects associated with *CEP250* or *CEP128* loss. Specifically, *CEP250* KO cells exhibit defects in centrosome cohesion, loss of the rootletin-CEP68 filaments, and impaired microtubule nucleation by centrosomes but also at the Golgi (Theile *et al*., 2023) (Fig EV5, Table 6). Thus, the strong migration phenotype observed in *CEP250* KO cells is likely attributable to increased centrosome distance and perhaps a decrease in microtubule nucleation activity but not the loss of the rootletin-CEP68 filaments. Microtubules anchored at subdistal appendages likely continue to contribute to cell migration in *CEP250* KO cells. This conclusion is supported by the behavior of *CEP128-250* cells, which had the strongest effect on cell migration of the analysed mutants. In contrast, *CEP128-68* cells behaved similarly to *CEP128* KO cells, consistent with the continued presence of centrosome cohesion and the limited role for the rootletin–CEP68 filaments in cell migration.

Taken together, we propose that centrosome cohesion mechanisms coordinate Golgi organization through multiple physical interactions and by regulating the distance between the two centrosomes. These interactions contribute differently to distinct cellular functions. Rootletin appears to play a more prominent role than CEP128 in maintaining the centrosome–Golgi connection and promoting Golgi compaction, whereas CEP128 has a stronger impact on cell migration (Table 6). Thus, although all centrosome cohesion mechanisms contribute to Golgi organization and cell migration, they appear to have distinct functional contributions to these processes.

## Materials and methods

- Key resources table
- Cell culture and transfection
- CRISPR-Cas9 genome editing
- Generation of stable cell lines
- Immunofluorescence staining
- Microtubule regrowth assay
- Scratch wound assay
- Golgi apparatus regrowth assay
- Super resolution microscopy (STED)
- Microscopy
- Live cell imaging
- Western blot analysis
- Immunoprecipitation
- Analysis and software

### Cell culture conditions and treatments

Human telomerase-immortalized retinal pigment epithelial (hTERT-RPE1), colorectal cancer (HCT116), and human embryonic kidney (HEK293T) cell lines were cultured in DMEM/F-12 (Gibco, Thermo Fisher). Osteosarcoma (U2OS) cells were maintained in DMEM/GlutaMAX™ (Thermo Fisher). For single-clone selection, all cell lines were transitioned to DMEM/F-12. All media were supplemented with 10% FBS, 1% L-glutamine, and 1% penicillin/streptomycin. Cells were maintained at 37°C with 5% CO_2_ and were routinely verified to be mycoplasma-free. Small interfering RNA (siRNA) was delivered using Lipofectamine RNAiMAX (Thermo Fisher Scientific) according to the manufacturer’s instructions. For plasmid DNA, RPE1 and HCT116 cells were transfected using Lipofectamine 2000 or Lipofectamine LTX (Thermo Fisher Scientific), while HEK293 cells were transfected using Polyethyleneimine (PEI). To achieve higher delivery efficiency for CRISPR-Cas9 and gRNA plasmids, RPE1 cells were electroporated using the Neon Transfection System (Thermo Fisher Scientific) following optimized manufacturer protocols. Doxycycline (Dox) was used for induction of gene expression using the TetON system (Clontech).For RPE1 cells TetON controlled genes were induced for 24 h using 1 µg/ml Dox. Cells were treated with 5 µM Nocodazole for 1 h at 37°C before fixation to depolymerize microtubules in the cells. Cells were incubated with EdU (from Click-iT EdU Cell Proliferation Kit for Imaging, Thermo Fischer) for labeling of S and G2 cells. Brefeldin A (BFA) was used to disperse the Golgi apparatus. Cells were treated with 2 µg/ml BFA for 2 hours at 37°C before fixation. The solvent DMSO was used for a mock control to Nocodazole and BFA treatment.

### CRISPR-Cas9 genome editing

CRISPR-Cas9 gene editing was performed following previously established protocols (Ran *et al*, 2013). gRNAs were designed and cloned into the pSpCas9(BB)-2A-GFP vector (PX458; Addgene, #48138). To ensure maximal gene inactivation, select knockouts were generated using a double cut strategy to delete large genomic fragments. 1.1□×□10^6^ RPE1 cells were electroporated with the two guide RNAs (6□µg per plasmid). GFP-positive cells were isolated by Fluorescence-Activated Cell Sorting (FACS). Single cells were then seeded into 96-well plates and cultured for 10-14 days to allow for colony formation. Resulting monoclonal clones were screened and confirmed through a multi-modal approach, including genomic PCR, sequencing, immunofluorescence (IF), and Western blotting (WB).

### EdU treatment to detect S-phase cells

To detect cells in S phase, cells were incubated with 10 nM EdU for 20 min before fixation, utilizing the Click-iT™ Plus EdU Alexa Fluor™ Imaging Kit (ThermoFisher, Cat. #C10638 and #C10640) according to the manufacturer’s protocol.

### Plasmids and constructs

pRetroX-Tet3G and pVSVG plasmids (Retro-X™ Tet-On® 3□G Inducible Expression System—Clontech) were used to generate RPE1 and HEK T293 cells with Tet-On® 3□G System. Dox-inducible tagged versions of different genes were generated by amplification of the cDNA via PCR and cloned into the pRetroX-TRE3G vector with NEBuilder® HiFi assembly mix. GalT-GFP construct was kindly gifted by Dr. Anne Schlaitz (BZH). AKAP450 cDNA was purchased from Genscript (<u>NM_005751.4</u>; OHu26045).

### Generation of stable cell lines

Stable cell lines with Dox-inducible expression were generated using the Retroviral Tet-On® 3G System (Takara Bio). Initially, RPE1 TetON parental cell lines were established according to the manufacturer’s protocol. Target genes were cloned into the pRetroX-TRE3G response vector. To produce the retrovirus, HEK293-GP packaging cells were co-transfected with the pRetroX-TRE3G construct and the pCMV-VSV-G envelope vector (Addgene #8454).

Target cells were then transduced by incubation with virus-containing medium. Following infection, stable genomic integration was ensured through antibiotic selection (Puromycin) or enriched by Fluorescence-Activated Cell Sorting (FACS). The resulting stable cell lines were then expanded for further characterization.

### Immunofluorescence staining

For indirect immunofluorescence (IF), cells were seeded on coverslips, washed once with PBS (except those after Nocodazole or BFA treatment) and fixed with cold methanol for 5 min at -20°C. The samples treated with BFA were directly fixed with cold methanol for 5 min at -20°C. Samples were permeabilized with 0.1% Triton X-100 for 10 min, unspecific binding was blocked with 10% (v/v) FBS for 1 hour followed by incubation with the primary antibodies in 3% (w/v) BSA for 1 hour in a humid chamber that was coated with aluminium foil. After three consecutive washing steps with PBS, the sample was incubated with the secondary antibody in 3% BSA for 30 min. DNA was co-stained with DAPI. The coverslip was mounted on a glass slide with Mowiol (Calbiochem) with and without PPD (p-phenylenediamine, 0.01%, anti-fading agent, Sigma) for conventional microscopy and with ProLong Gold Antifade Mountant or ProLong Glass Antifade Mountant (ThermoFischer) for STED microscopy. For STED microscopy, cells were seeded on 12 mm precision coverslips #1.5H (Marienfeld 0117520) and fixed with cold methanol for 5 min at -20°C. Samples were permeabilized with 0.1% Triton X-100 for 10 min, unspecific binding was blocked with 10% (v/v) FBS overnight. Primary antibodies were incubated in 3% (w/v) BSA for 1 hour in a humid chamber. After three consecutive washing steps with PBS (twice 5 min and once 30 min), the sample was incubated with the secondary antibody in 3% BSA for 1 hour. Click-iTTM Plus EdU Alexa FluorTM 555 Imaging Kit (Thermo Fischer) was used to specify S and G2 cells. The EdU staining was performed after blocking and before the primary antibody incubation with three times washing steps with PBS in between.

### Golgi reformation assay

RPE 1 cells were seeded 24 hours prior to the drug treatment. Cells were treated with either DMSO or 2 µg/ml of Brefeldin A (Misumi *et al*., 1986) diluted in DMEM F-12 complete media for 2 hours at 37°C. 2-hour drug timepoint cells were fixed using ice cold methanol for 5 min. For the washout samples, cells were washed two times with PBS and complete DMEM F-12 was added to the samples and incubated at 37°C. Cells were then fixed after 30 min and 120 min time points. Immunofluorescence was conducted as mentioned before.

### Scratch wound assay

Cells were seeded in an Incucyte^®^ Imagelock 96-Well plate. Scratch wound Incucyte^®^ Woundmaker Tool was used to generate the scratch on all the samples simultaneously. The plate was then imaged using the Incucyte ® Live-Cell Analysis System at 10x objective with 1-hour intervals. Data analysis and raw images were processed using the included scratch wound assay Incucyt software.

### Microtubule regrowth assay

For the cold treatment, to depolymerize MT from the centrosomes, cells were put on ice and incubated at 4°C for 40 min. Afterwards, cells were washed with 1X PBS at 37°C and placed on a pre-heated metal rack (at 37°C) for 5 min to allow microtubule re-polymerization. Before fixation with cold methanol (−20°C) and IF staining, washed out with 1X PBS.

### Microscopy

The DeltaVision RT imaging system (GE Healthcare), Incucyte® live-cell imaging system (Sartorius), and STED microscope (Abberior Expert Line) were used in this study. For conventional imaging of fixed samples and subsequent quantitative analysis, the DeltaVision RT were employed. Images acquired via DeltaVision were processed (deconvolution and/or projection) using softWoRx software and quantified in Fiji. Live cell imaging experiments were conducted using the DeltaVision Rt system and the Incucyte ® Live-Cell Analysis System. All sample preparation and imaging procedures were performed according to the manufacturers’ standardized protocols.STED measurements were performed on an Abberior Expert Line STED microscope using a 100× UPlanSApo oil immersion objective (Olympus). For STED imaging, a 40 MHz pulsed 775 nm depletion laser (3 W at laser head) and pulsed excitation lasers of 640 nm (1.2 mW at laser head) and 561 nm (203 µW at laser head) were used. STED images were taken of a 4 µm × 4 µm field of view with 20 nm pixels, using a 0.7 AU pinhole, and gating with 10 ns width. Dual color STED images were made by imaging both channels in separate line steps, using detection windows of 575-630 nm for 561 nm excitation (for STAR 580) and 650-754 nm for 640 nm excitation (for STAR 635P and STAR RED). For STED imaging of rootletin (STAR 580), we used 25-30% 561 nm excitation power, 100% STED power, and 60 µs total pixel dwell time over 4-line repetitions. For STED imaging of AKAP450 (STAR 635P), we used 2.5% 640 nm excitation power, 30% STED power, and 45 µs total pixel dwell time over 3-line repetitions. For STED imaging of AKAP450^PACT^-HA and PCNT^PACT^-HA (STAR RED), we used 4% 640 nm excitation power, 25% STED power, and 90 µs total pixel dwell time over 6-line repetitions.

### Live cell imaging

For live cell imaging, cells were grown in Ibidi µ-Slide 8-well plates (Ibidi) and labelled with SPY650-DNA dye at a dilution of 1:7000 for 2 hours before imaging. For BFA washout experiments, 2 µg/ml of BFA was added 2 hours before imaging Golgi reassembly.5µM Nocodazole was added 1 hour before imaging for centrosome separation. Images were acquired (DeltaVision RT system) as z-stacks with a 60x objective every 2□min for 2 hours at 37°C. Maximum intensity projections were created using SoftWoRx. Intercentrosomal distance and centrosome-Golgi distance were measured using FIJI. 15 µm stack images were taken with 1 µm interspace between each stack. For all live cell experiments, HEPES-buffered medium without phenol red was used.

### Electron Microscopy

Non transformed RPE1 cells were seeded on coverslips and cultured at 37°C and 5% CO_2_ till they reached a confluency of approximately 60-80%. Cells were rinsed twice with phosphate buffer (PBS) and pre-fixed with a mixture of 2.5% GA/1.6% PFA/2% sucrose in 50 mM cacodylate buffer for 30 min at room temperature. After washing of cells with cacodylate buffer several times, cells were post-fixed with 2% OsO_4_ for approximal 45 min on ice, in darkness. Cells were washed intensively with dH_2_O and incubated overnight at 4°C in 0.5% uranyl acetate/water. On the following day coverslips were rinsed again with dH_2_O and subsequently stepwise dehydrated with ethanol (40-60-70-80-90-95-100% EtOH). Coverslips were immediately placed on capsules filled with Spurr-resin (Sigma-Aldrich) and polymerized at 60°C for approximately two days. In resin embedded cells were serial-sectioned using a Reichert Ultracut S Microtome (Leica Instruments, Vienna, Austria) to a thickness of 80 nm. Sections of cells were post-stained with lead citrate and imaged at a Jeol JE-1400 (Jeol Ltd., Tokyo, Japan), operating at 80 kV, equipped with a 4k x 4k digital camera (F416, TVIPS, Gauting, Germany). Micrographs were adjusted in brightness and contrast applying Fiji software.

### Immunoprecipitation

To test subdomains of AKAP450 interaction with CEP68, *AKAP450-FLAG* constructs were co-expressed with *CEP68-HA.* HEK293T cells were transfected with PEI, induced with doxycycline, and harvested 24 hours post-transfection. Cells were lysed in buffer [10 mM Tris-HCl (pH 7.5), 150 mM NaCl, 0.5 mM EDTA, 0.5% NP-40, 1 mM PMSF, 10 U/µl Benzonase, and Roche complete protease inhibitor cocktail. Lysates were incubated with HA conjugated magnetic beads for 1 hour at 4°C under rotation. After three washes with a high-salt buffer (10 mM Tris-HCl pH 7.5, 300 mM NaCl), beads were directly boiled with 30 µl of 4x Laemmli buffer.

### Western blot analysis

Protein lysates were prepared by washing cells once with PBS followed by lysis in 2x Laemmli buffer on ice. Cells were harvested using a cell scraper, collected into 1.5 ml tubes, and briefly centrifuged to eliminate air bubbles. To reduce viscosity, benzonase (Merck, 1:1000) was added, and samples were incubated on ice for 10 min. Lysates were clarified by centrifugation at 16,000 rpm for 15 min at 4°C. The resulting supernatant was transferred to fresh tubes and denatured at 95°C for 5 min. Proteins were resolved by SDS-PAGE and transferred onto methanol-activated PVDF membranes. Membranes were blocked in 5% skimmed milk in TBST (TBS, 0.1% Tween 20) for 30 min at room temperature. Following three 5 min washes in TBST, membranes were incubated with primary antibodies (diluted in 3% BSA in TBST) overnight at 4°C. After further washing, membranes were incubated with secondary antibodies (diluted in 5% skimmed milk in TBST) for 30 min at room temperature. Protein bands were visualized using Clarity or Clarity Max ECL substrates (Bio-Rad) and imaged with a Fujifilm LAS-4000 system.

### Image Analysis and Radial Quantification

Cells were segmented from microscopy images using Cellpose (Pachitariu *et al*., 2025), excluding border-touching cells and objects smaller than 500 pixels in area. Following median background subtraction of non-cellular signal, the Golgi radial dispersion was quantified relative to the intensity-weighted center of mass of either the Golgi or the centrosome. Specifically, radial dispersion of each cell was characterized by the 70% intensity quantile, defined as the radius covering 70% of the total Golgi signal. Outlier cells were removed within each experimental condition using a 1.5 x interquartile range (IQR) criterion on the Q70 values; the Q70 average over the remaining cells is shown in the figures.Custom Python pipelines are publicly available on GitHub: (https://github.com/anders-biostat/golgi-radial).

### Data Analysis

All STED images were processed using Wiener deconvolution (<u>10.5281/zenodo.17600326</u>). All raw data is available on DataverseNL. For STED images, line profiles were drawn in Fiji using a segmented line with spline fit and a line width of 8 pixels.

## Supporting information

Expanded View EV1-EV5

## Acknowledgement

We thank Dr. Cornelia Sala for helpful discussion. We thank Ursula Jäkle for support and technical assistance. We are thankful to Dr. H. Lorenz and Christian Hoerth of the ZMBH Imaging Facility for their assistance with the scratch wound analysis. Additionally, we acknowledge Dr. M. Langlotz from the ZMBH FACS Facility for their support in sorting GFP-positive Cas9 transfected cells, GalT-GFP stable cell lines and mRuby2 stable cell lines. We thank Dr. Anne Schlaitz from the Heidelberg University Biochemistry Center (BZH) for sharing the GalT-GFP construct. The *AKAP450* KO RPE1 cell line was obtained from Dr. Rosa Rios (Gavilan *et al*., 2018). We thank Gislene Periera for the ODF2 antibody.

This project was funded by the Deutsche Forschungsgemeinschaft: Schi 295/6-3 to ES and OCENW.M #21.106 to R.V.

## Data availability

The data that support the findings of this study and the plasmids generated in this study are available from the corresponding author Elmar Schiebel upon reasonable request.

## Author Contributions

**Elmar Schiebel:** Conceptualization; data curation; funding acquisition; writing – original draft; project administration; writing – review and editing. **Aadyasha Mishra:** Conceptualization; validation; investigation; visualization; methodology; writing – original draft; writing – review and editing. **Sietse J Dijt:** Investigation; methodology; writing-review and editing. **Laura Theile:** Conceptualization; validation. **Annett Neuner:** Investigation; methodology. **Tobias Benedikt Roth:** Software; investigation; methodology. **Simon Anders:** Software; investigation; methodology. **Rifka Vlijm:** Supervising STED project, discussions.

## Expanded View figure legend

**Expanded View Figure 1 Ninein centrosomal localization depends on CEP250 and CEP128 and Rootletin and CEP68 levels are unaffected by AKAP450 loss.**

**A** EM images of centrioles from WT *and NIN* KO cells. The knockout cells lines have have a slight defect in the microtubule wall (as seen in the longitudinal view, pinkarrows) with the subdistal appendages being formed. Scale bars: 200Onm. Abbreviations: DA, distal appendages; SDA, subdistal appendages.

**B** Representative fluorescent image of RPE1 WT, *CEP250* KO, *CROCC* KO, *CEP68* KO, *CEP128* KO, *CEP128-250* KO, *CEP128-68* KO cells. Ninein (red) was stained and γ-tubulin (green) was used to mark centrosomes and DNA was stained with DAPI. The enlargements at the top show the centrosome signal of the cell in the main panel. White scale bars, 5 µm. Yellow scale bars, 1 µm.

**C** Quantification of the normalized ninein signal intensities (a.u.) at the centrosome in (A). n>100 cells per experiment, N=3 independent experiments. Bar and error represent mean and SD. Unpaired *t*-test.

**D** Representative fluorescent image of RPE1 WT and doxycycline induced *NIN* KO + *NIN*-mNG rescue cell lines. Rootletin (red) and CEP68 (red) was stained γ-tubulin (yellow) was used to mark centrosomes, *NIN*-mNG (green) is expressed with 2 ng doxycycline and DNA was stained with DAPI. The enlargements at the top show the centrosome signal of the cell in the main panel. White scale bars, 5 µm. Yellow scale bars, 1 µm.

**E, F** Quantification of the rootletin (E) and CEP68 (F) signal intensities (a.u.) at the centrosome in (D). n>100 cells per experiment, N=2 independent experiments. For *NIN* KO cells see Fig 2B and C. Bar and error represent mean and SD. Unpaired *t*-test.

**G** Representative fluorescent image of RPE1 WT and *AKAP450* KO cells. Rootletin (red) and CEP68 (red) was stained γ-tubulin (green) was used to mark centrosomes and DNA was stained with DAPI. The enlargements at the top show the centrosome signal of the cell in the main panel. White scale bars, 5 µm. Yellow scale bars, 1 µm. **H, I** Quantification of the rootletin (H) and CEP68 (I) signal intensities (a.u.) at the centrosome in (G). n>100 cells per experiment, N=2 independent experiments. Bar and error represent mean and SD. Unpaired *t*-test.

**Expanded View Figure 2 AKAP450 plays a role in interphase centrosome cohesion in U2OS and HCT116 but not RPE1 cells.**

**A, C, F** Fluorescent images of centrosomes in RPE1, U2OS, HCT116 cells under siRNA depletion of non-specific siRNA control (siControl), or C-Nap1, rootletin and AKAP450 siRNA, without and with microtubule depolymerization using 5 µM Nocodazole for 1 h. γ-tubulin (green) and GM130 (red) was used as a centrosome marker and DNA was stained with DAPI. The boxes on top show enlargement of the centrosome signals of the cell in the main panel. White scale bars, 5 µm. Yellow scale bars, 1 µm.

**B, D, F** Quantification of cells with separated centrosomes (centrosome distance >2 µm for RPE1, U2OS cells and >1 µm for HCT116 cells) from RPE1 WT (A), U2OS WT (C) and HCT116 WT (E) cells. n=50 cells per experiment, N=3 independent experiments. Bars and errors represent mean and SD. Unpaired *t*-tests.

**Expanded View Figure 3 Spatial relationship of the centrosomes with Golgi in G1 cells.**

**A** Representative fluorescent image of RPE1 WT, *CEP250* KO, *CROCC* KO, *CEP68* KO, *CEP128* KO, *CEP128-250* KO, *CEP128-68* KO cells. ODF2 (magenta) was stained as a mother centriole marker, γ-tubulin (green) was used to mark centrosomes, GM130 was used as a Golgi marker (red) and DNA was stained with DAPI. White scale bars, 5 µm.

**B** Quantification of centrosomes position in respect to the Golgi. Percentage of cells in the WT and knockout cell lines showing attachment of the centrosome to the Golgi (A). Bars and errors represent mean and SD. n>30 cells, N=3 per cell line for each experiment.

**Expanded View Figure 4 Cell cycle progression only mildly influences Golgi organization.**

**A** Representative fluorescent image of RPE1 WT and *CEP250* KO cells. EdU staining (yellow) was implemented to identify S phase cells, γ-tubulin (green) was used to mark centrosomes, GM130 (red) was used as a Golgi marker and DNA was stained with DAPI. The enlargements at the top show the centrosome signal of the cell in the main panel. White scale bars, 5 µm. Yellow scale bars, 1 µm.

**B** Percentage of cells in the WT and *CEP250* KO showing the mentioned Golgi phenotype in G1, S and G2 cells in (A). n≥30 cells, N=2 per cell line for each experiment.

**C** Quantification of Golgi morphology phenotypes in interphase G1/S RPE1 WT, *CEP250* KO, *CROCC* KO, *CEP68* KO, *CEP128* KO cells. Representative fluorescent images indicate the different phenotypes. γ-tubulin (green) was used to mark centrosomes, GM130 (red) was used as a Golgi marker. Bar graph showing the percentage of interphase cells displaying (compact/half-moon/extended) Golgi morphology across the indicated cell lines. Cells were classified based on GM130. CENPF staining (not shown in the figure) was used to distinguish the G2 cells. n≥30 cells per experiment, N=3 independent experiments. Bar and error bars represent mean and SD.

**D** Representative fluorescent images of RPE 1 *CEP128-250* KO, *CEP128-68* KO, *CEP250* KO+*CEP250* rescue cell line. γ-tubulin (green) and GM130 (red) antibodies were used for indirect fluorescent analysis. The cells were imaged at three different timepoints; two hours after drug treatment, 30 minutes after washout and 120 minutes after washout. DMSO treated cells were used as control. DNA was stained with DAPI. Scale bars: 5 µm.

**E** Radius of a circle was drawn around the centre of mass of the cell’s Golgi signal (GM130 signal) such that the circle encloses 70% of the total signal intensity of the GM130 signal. WT and *CEP250* KO are as in Fig 6D. N=3 independent experiments; n≥80 cells per cell line. Faster accumulation (smaller diameter) means more dense distribution.

**F** Quantification of centrosomes position in respect to the Golgi after 2 h washout of BFA from cells in Fig 6B. Percentage of cells in the WT and knockout cell lines showing attachment of the centrosome to the Golgi. Bars and errors represent mean and SD. n≥30 cells N=2 per cell line for each experiment.

**Expanded View Figure 5 Disrupting centrosome cohesion alters microtubule regrowth and reduces microtubule acetylation.**

**A** Experiment design for microtubule regrowth assay. Cells were incubated at 4°C for 1 h to allow depolymerization of microtubules and then incubated at 37°C for 5 minutes to allow repolymerization of microtubules.

**B** Representative fluorescent image of RPE1 WT, *CEP250* KO, *CROCC* KO, *CEP68* KO and *CEP128* KO cells. Acetylated alpha-tubulin (grey/green) was stained as a stable microtubule marker, γ-tubulin (yellow) was used to mark centrosomes, GM130 (red) was used as a Golgi marker and DNA was stained with DAPI. Scale bars, 5 µm.

**C** Quantification of mean acetylated tubulin signal intensities (a.u) at the centrosome and Golgi in (A). n>100 per experiment, N=2 independent experiments. Bar and error represent mean and SD. Unpaired *t*-test.

**D** Quantification of centrosomes position in respect to the Golgi after 5 minutes of microtubule regrowth. Percentage of cells in the WT and knockout cell lines showing attachment of the centrosome to the Golgi (Fig 6A). Bars and errors represent mean and SD. n>50 cells per cell line for each experiment, N=3.

**E** Representative image of scratch wound analysis of RPE1 *CEP250* KO + *CEP250* rescue cell line and *AKAP450* KO as a positive control of the migration defect. Phase contrast overlaid images illustrate wound gaps at 0 h, 8 h and 18 h. WT and *CEP250* KO curve are the same as in Fig 7B. Scale bars: 600 µm.

**F** Relative wound closure of the indicated cell lines over 24 h, measured using the Incucyte Scratch Wound assay. Wound closure is expressed as relative wound density (%). Images were acquired every 2 h with a 10× objective. WT and *CEP250* KO are as in Fig 7B.

## Appendix Figures

**Appendix Figure S1 Localization of AKAP450 in mitotic cells in RPE1, U2OS and HCT116 cells**

**A** Representative line-scan intensity profile of centrosome showing the centrosomal distribution of AKAP450 (green) and γ-tubulin (magenta) in U2OS cells (Fig 1B). Fluorescence intensity (arbitrary units, a.u.) is plotted as a function of distance (µm) along the x axis. n=15 cells pooled from N=3 independent experiments were analyzed.

**B** Representative line-scan intensity profile of centrosome showing the centrosomal distribution of AKAP450 (green) and γ-tubulin (magenta) in HCT116 cells (Fig 1B). Fluorescence intensity (arbitrary units, a.u.) is plotted as a function of distance (µm) along the x axis. n=15 cells pooled from N=3 independent experiments were analyzed.

**C, D and E** Representative fluorescent images of RPE1 WT, U2OS WT and HCT116 WT mitotic cells. AKAP450 (green) was stained, PCNT (red) and γ-tubulin (magenta) was used to mark the spindle poles. Scale bars, 5 µm.

**F** Quantification of AKAP450 signal intensities (a.u) at the centrosome in interphase and mitotic cells (D, E, F). n>30 per experiment, N=2 independent experiments. Bar and error represent mean and SD. Unpaired *t*-test.

**Appendix Figure S2 Localization of AKAP450 in RPE1, U2OS and HCT116 using confocal and STED images.**

**A** Confocal γ-tubulin (blue) images corresponding to the STED images of rootletin (green) and AKAP450 (red) presented in Fig 1. Scale bars, 1 µm.

**B** STED images of rootletin (green) and AKAP450 (red) in RPE1, HCT116 and U2OS cells, showing rootletin fibers that are disconnected from the main rootletin network around the centrosome. Such disconnected rootletin fibers were found in a small amount (6-7%) of STED images of centrosomes. In total, 124 (RPE1), 178 (HCT116) or 127 (U2OS) randomly selected centrosomes from 3 samples per cell type were imaged and classified. For such rootletin fibers, AKAP450 was present around the fiber. White scale bars, 1 µm yellow scale bars, 200 nm.

**C** Confocal overviews and STED images of rootletin, AKAP450 and confocal images of GalT-GFP as seen in Fig1 H, showing the different centrosome-Golgi situations in RPE1 GalT-GFP WT cells. The white box in the confocal overview shows the location of the STED image. In total, 107 randomly selected centrosomes from 3 samples were imaged and classified. Typically, centrosomes were found at the Golgi, and in some cases, AKAP450 appeared to extend from the rootletin fibers into the Golgi. The images on top are identical to the images shown in Fig 1H, bottom. White scale bars, 2 µm. Yellow scale bars, 1 µm.

**D** STED images of AKAP450 (red), rootletin (green), and confocal images of GalT-GFP (gray) at centrosomes in RPE1 GalT-GFP cells, showing examples of rootletin fibers going along or towards GalT-GFP. White scale bars, 1 µm. Yellow scale bars, 500 nm.

**E** Confocal overviews and STED images of rootletin (green), AKAP450 (red) and confocal images of GalT-GFP (blue) in RPE1 GalT-GFP *CEP250* KO cells. The white box in the confocal overview shows the location of the STED image. In total, confocal overviews of 389 GalT-GFP-positive cells from 3 samples were made and classified. Most cells had similar Rootletin fibers and these fibers were nearly always at the Golgi. White scale bars, 10 µm. Yellow scale bars, 1 µm.

**F** STED images of AKAP450 (red), rootletin (green), and confocal images of GalT-GFP (gray) at centrosomes in RPE1 GalT-GFP *CEP250* KO cells, showing there are still Rootletin fibers in the Golgi. White scale bars, 1 µm. Yellow scale bars, 500 nm.

**Appendix Figure S3 Generation and characterization of RPE1 *CEP128* KO and effect of *CEP128* in centrosome cohesion.**

**A** Scheme of *CEP128* gene with the selected gRNA2 and gRNA4 used for the of deletion in RPE1 *CEP128* KO. The sgRNAs used to generate the knockout targeting exon 3 and intron 8 are shown, resulting in a deletion of 12.9 kb. The gene structure was drawn using FancyGENE (Rambaldi & Ciccarelli, 2009).

**B** Scheme of the *CEP128* WT gene and the domain architecture of the protein.

**C** Chromatogram of the sequenced knockout clone showing successful deletion. The clone shows a homozygous deletion, in which both alleles have the large deletion following exon 3, after which the sequence of intron 8 is following. In comparison, in the *CEP128* WT allele of control cells, the sequence of intron 3 can be amplified with PCR and subsequently sequenced.

**D** Immunoblot of endogenous CEP128 in RPE1 WT and *CEP128* KO #20, *CEP128-250* KO #57, *CEP128-68* KO #109 cells. GAPDH was used as loading control.

**E** Fluorescent images of CEP128 (red) and γ-tubulin (green) in the indicated RPE1 cells. DNA was stained with DAPI. The insets show enlargement of the centrosome signals of the cell in the main panel. Scale bars in white: 5 µm; scale bars in yellow: 1 µm.

**F** Quantification of normalised relative CEP128 intensity at the centrosome in RPE1 cells from (E). n=50 cells, N=2 independent experiments. Bar and error represent mean and SD. Unpaired *t*-test.

**G** Quantification of cells with separated centrosomes (centrosome distance >2□µm) from RPE1 samples with DMSO as control and 5 µm of Nocodazole. n=□50 cells per experiment, N=2 independent experiments. Bar and error represent mean and SD. Unpaired *t*□test.

**H** EM images of centrioles from WT *and CEP128* KO *cells*. WT cells show clear distal appendages (DA; purple arrows) and subdistal appendages (SDA; pink arrows). The *CEP128* KO cells have loss of subdistal appendages (as seen in the longitudinal view) with the distal appendages still intact (purple arrow). Mother centrioles were identified in *CEP128* KO by checking centrioles presenting the ciliary shaft (blue circles) and ciliary vesicles (blue stars) and Scale bars: 200Onm.

**Appendix Figure S4 Localisation of the PACT domain in confocal and STED microscopy.**

**A** Confocal γ-tubulin images corresponding to the STED images of Rootletin (green) and AKAP450^PACT^-HA (red) in the AKAP450^PACT^-HA cell line presented in Fig 3. AKAP450^PACT^-HA is present at the centrosomes, marked by γ-tubulin (blue).

**B** Confocal γ-tubulin images corresponding to the STED images of Rootletin (green) and PCNT^PACT^-HA (red) in the PCNT^PACT^-HA cell line presented in Fig 3. PCNT^PACT^-HA is present at the centrosomes, marked by γ-tubulin.

**(A-B)** Scale Bars, 1 µm.

**Appendix Figure S5 STED images of Rootletin and AKAP450^PACT^-HA.**

**A** Different Rootletin (green)/AKAP450^PACT^-HA (red) organizations in RPE1 AKAP450^PACT^-HA cells. In total, n=90 randomly selected centrosomes from N=3 samples were imaged and classified.

**B** Different centrosome separations were found in RPE1 *AKAP450^PACT^-HA CEP250* KO cells. Typically, no Rootletin fibers were found near the centrosomes.

**C** STED images of Rootletin (green) and AKAP450^PACT^-HA (red) in RPE1 *AKAP450^PACT^-HA CEP250* KO cells, showing AKAP450^PACT^-HA was typically not present on these detached Rootletin fibers.

**D** Typical Rootletin (green)/AKAP450^PACT^-HA (red) organization in RPE1 *AKAP450^PACT^-HA CEP68* KO cells.

**(A-D)** Scale Bars, 1 µm.

**Appendix Figure S6 STED images of Rootletin and PCNT^PACT^-HA show similar organizations as for AKAP450^PACT^-HA.**

**A** Different Rootletin (green)/PCNT^PACT^-HA (red) organizations in RPE1 *PCNT^PACT^-HA* cells. In total, n=90 randomly selected centrosomes from N=3 samples were imaged and classified.

**B, C** Different centrosome separations were found in RPE1 *PCNT^PACT^-HA CEP250* KO cells. Typically, no Rootletin fibers were found near the centrosomes.

**D** Typical Rootletin (green)/PCNT^PACT^-HA (red) organization in RPE1 *PCNT^PACT^-HA CEP68* KO cells.

**(A-D)** Scale Bars, 1 µm.

**Appendix Figure S7 Confirmation of siRNA depletion.**

**A, B, C** Depletion efficiency of the indicated siRNAs in RPE1 WT, U2OS WT and HCT116. Quantification of the relative intensity at the centrosome by IF. N>30 cells; a representative dataset from 2 independent experiments is shown. Dots represent the relative intensity of the protein at the centrosome in protein at the centrosome in each cell. Lines represent mean; error bars represent SD. Unpaired *t*-test.

**Appendix Figure S8 Centrosome linker mutants affect total microtubule regrowth.**

**A** Representative fluorescent image of RPE1 WT, *CEP250* KO, *CROCC* KO, *CEP68* KO, *CEP128* KO, *CEP128-250* KO, *CEP128-68* KO cells. α-tubulin was stained to mark the microtubules (black), GM130 was used as a Golgi marker (red). Scale bars, 5 µm. Blue line shows contour of the cell.

**B** Quantification of cells from (A). n>50 cells, N=2 independent experiments. Mean intensity of the total α-tubulin signal in the cell. Bar and error represent mean and SD. Unpaired *t*-test.

## Movie EV1-EV4 Legend

**Movie EV1. Example movie of RPE1 WT with siControl or si AKAP450 with DMSO and NOCO treatment.**

Related to Figure 4. Time-lapse movies (5 min intervals) of RPE1 WT siControl DMSO, RPE1 WT siAKAP450 DMSO, RPE1 WT siControl NOCO and RPE1 WT siAKAP450 NOCO cells expressing γ-tubulin-mRuby2 (red) as marker for the centrosomes. The DNA was marked with SPY-DNA650 (grey). Cells were incubated with DMSO as solvent control for the Nocodazole samples in Movie EV1.

**Movie EV2. Example movie of HCT116 WT with siControl or si AKAP450 with DMSO and NOCO treatment.**

Related to Figure 4. Time-lapse movies (5 min intervals) of HCT116 WT siControl DMSO, HCT116 WT siAKAP450 DMSO, HCT116 WT siControl NOCO and HCT116 WT siAKAP450 NOCO cells expressing γ-tubulin-mRuby2 (red) as marker for the centrosomes. The DNA was marked with SPY-DNA650 (grey). Cells were incubated with DMSO as solvent control for the Nocodazole samples in Movie EV2.

**Movie EV3. Example movie of centrosome-Golgi dynamics in RPE1 WT, *CEP250* KO, *CROCC* KO and *CEP128* KO.**

Related to Figure 5. Time-lapse movies (5 min intervals) of RPE1 WT, *CEP250* KO, *CROCC* KO and *CEP128* KO cells expressing γ-tubulin-mRuby2 (red) and GalT-GFP (green) as markers for the centrosomes and Golgi, respectively. The DNA was marked with SPY-DNA650 (magenta).

**Movie EV4. Example movie of Golgi reformation in RPE1 WT, *CEP250* KO, *CROCC* KO and *CEP128* KO.**

Related to Figure 6. Time-lapse movies (5 min intervals) of RPE1 WT, *CEP250* KO, *CROCC* KO and *CEP128* KO cells expressing γ-tubulin-mRuby2 (red) and GalT-GFP (green) as markers for the centrosomes and Golgi, respectively. The DNA was marked with SPY-DNA650 (magenta).

