## Supplementary figures and images for "Control of Golgi Architecture Through Three Centrosome Cohesion Pathways"

### Expanded View EV1-EV5

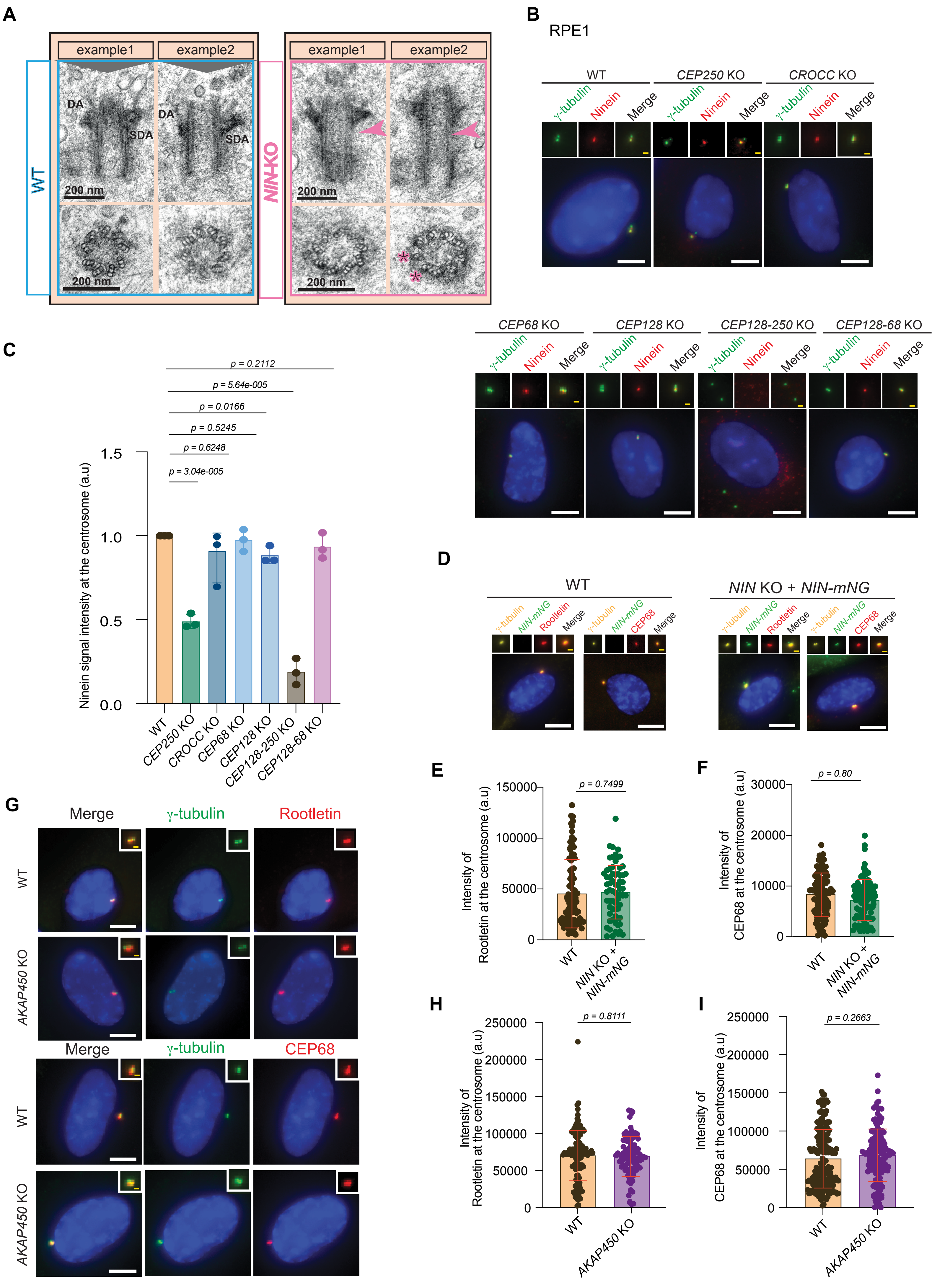

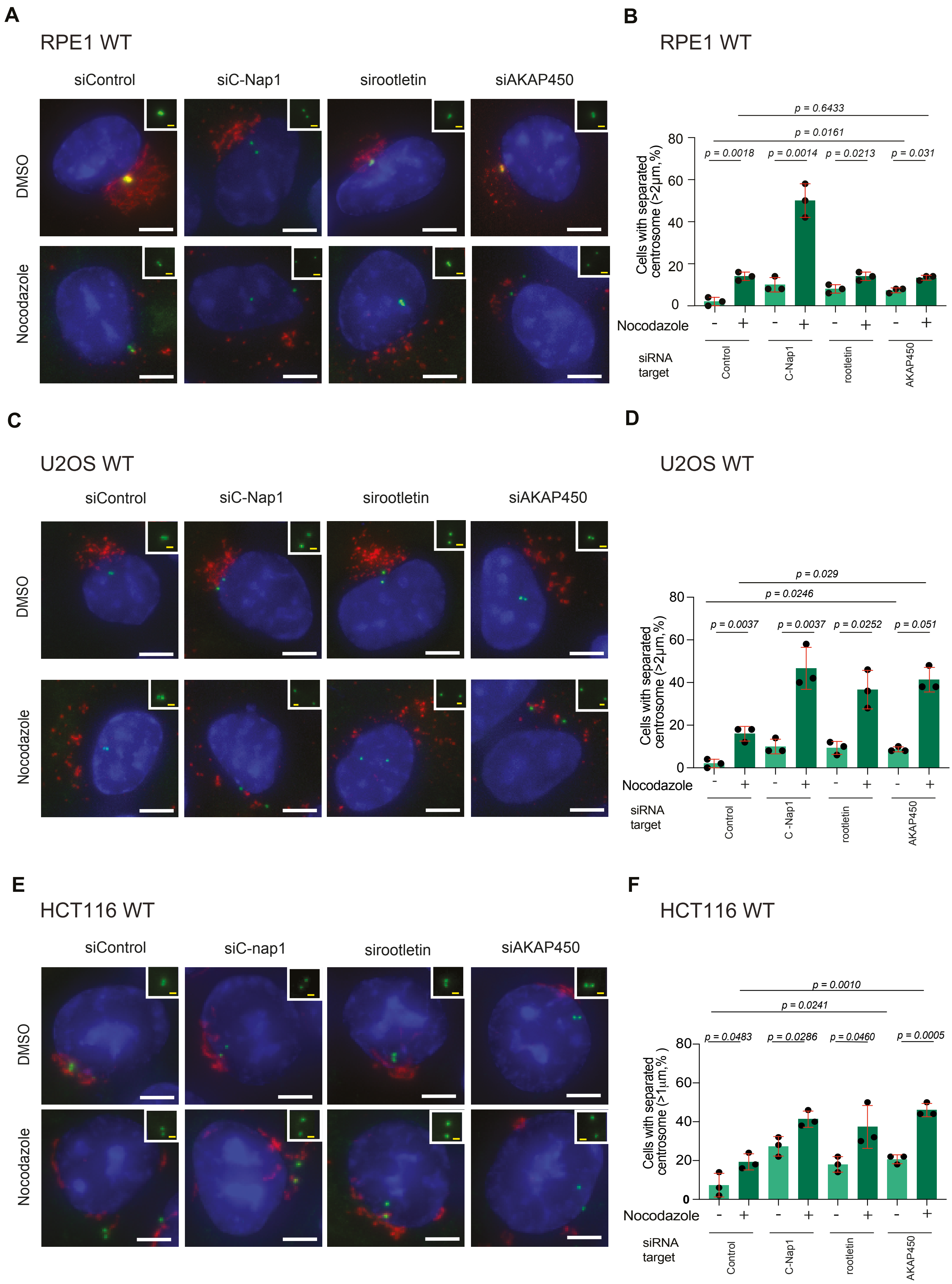

**Figure EV3**

**A**

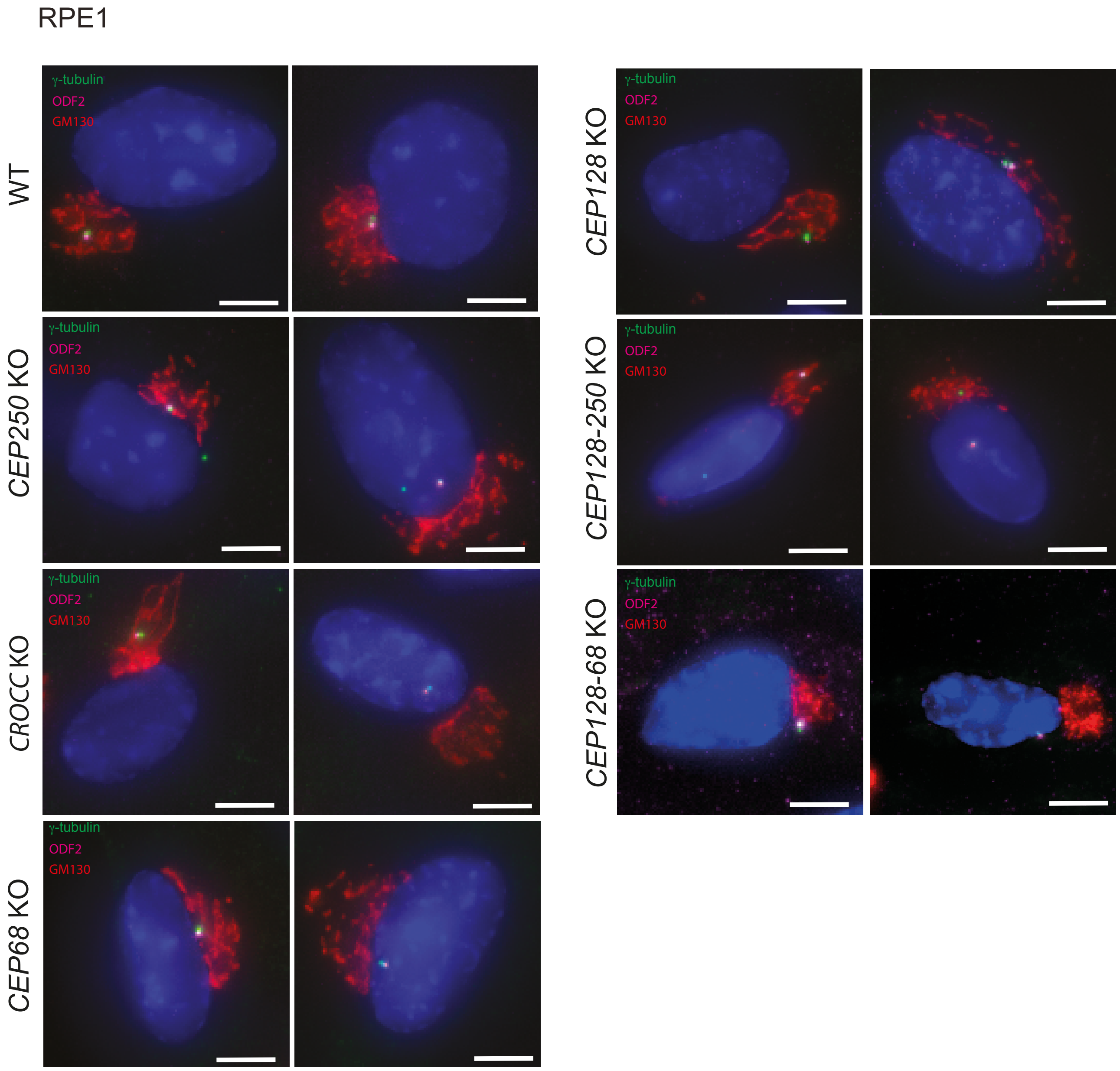


**B**

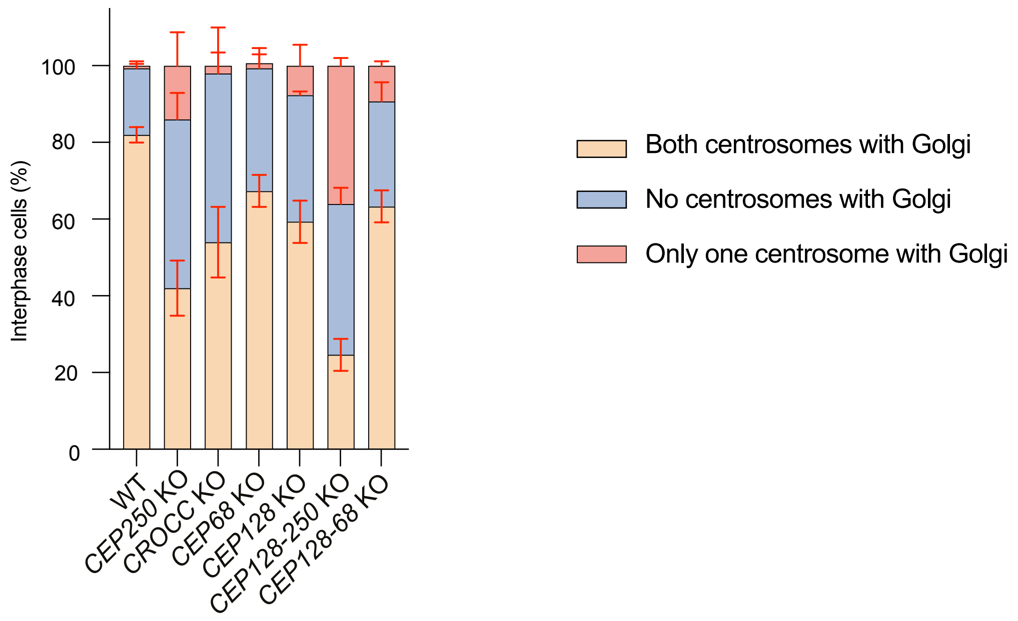

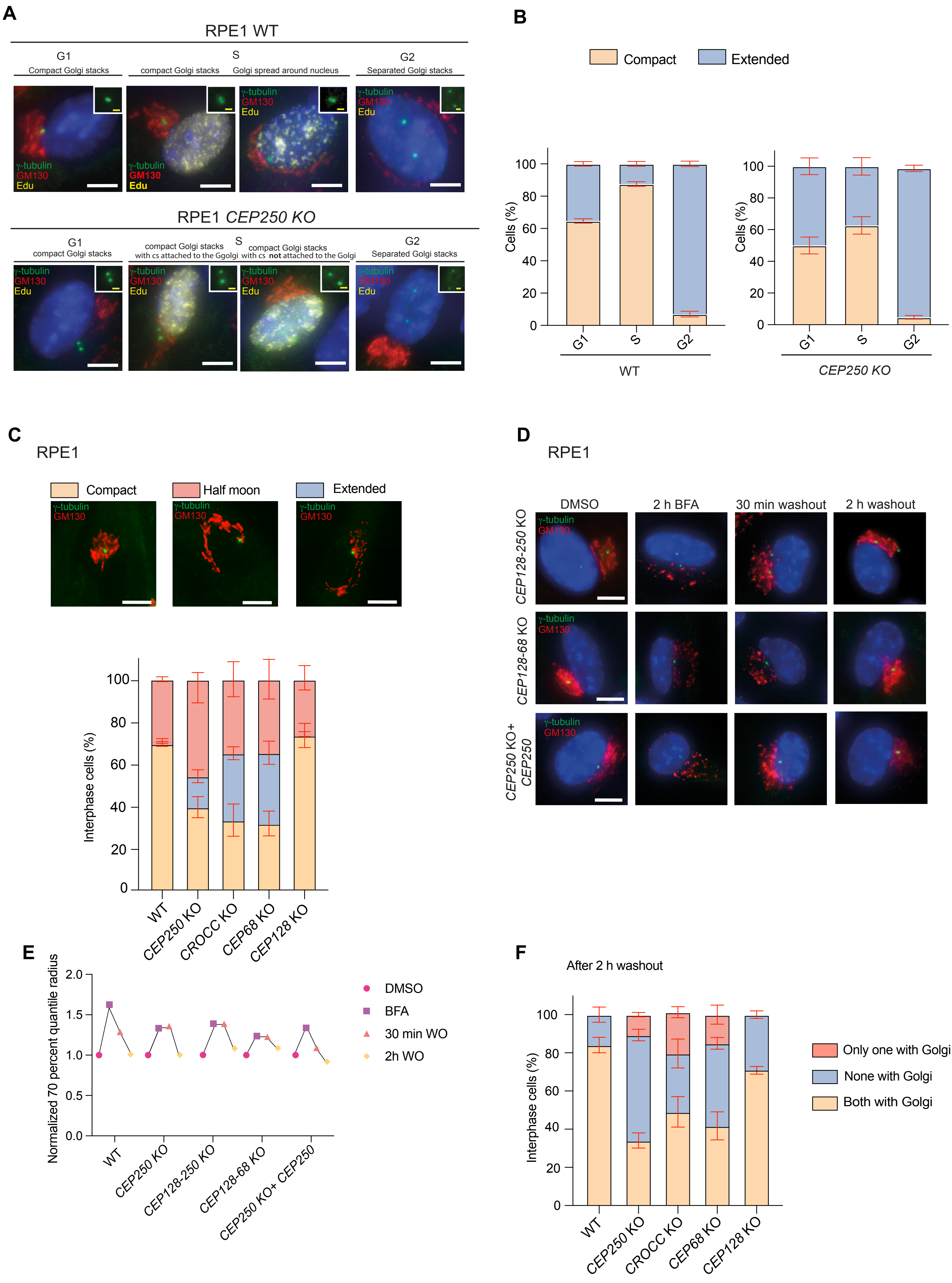

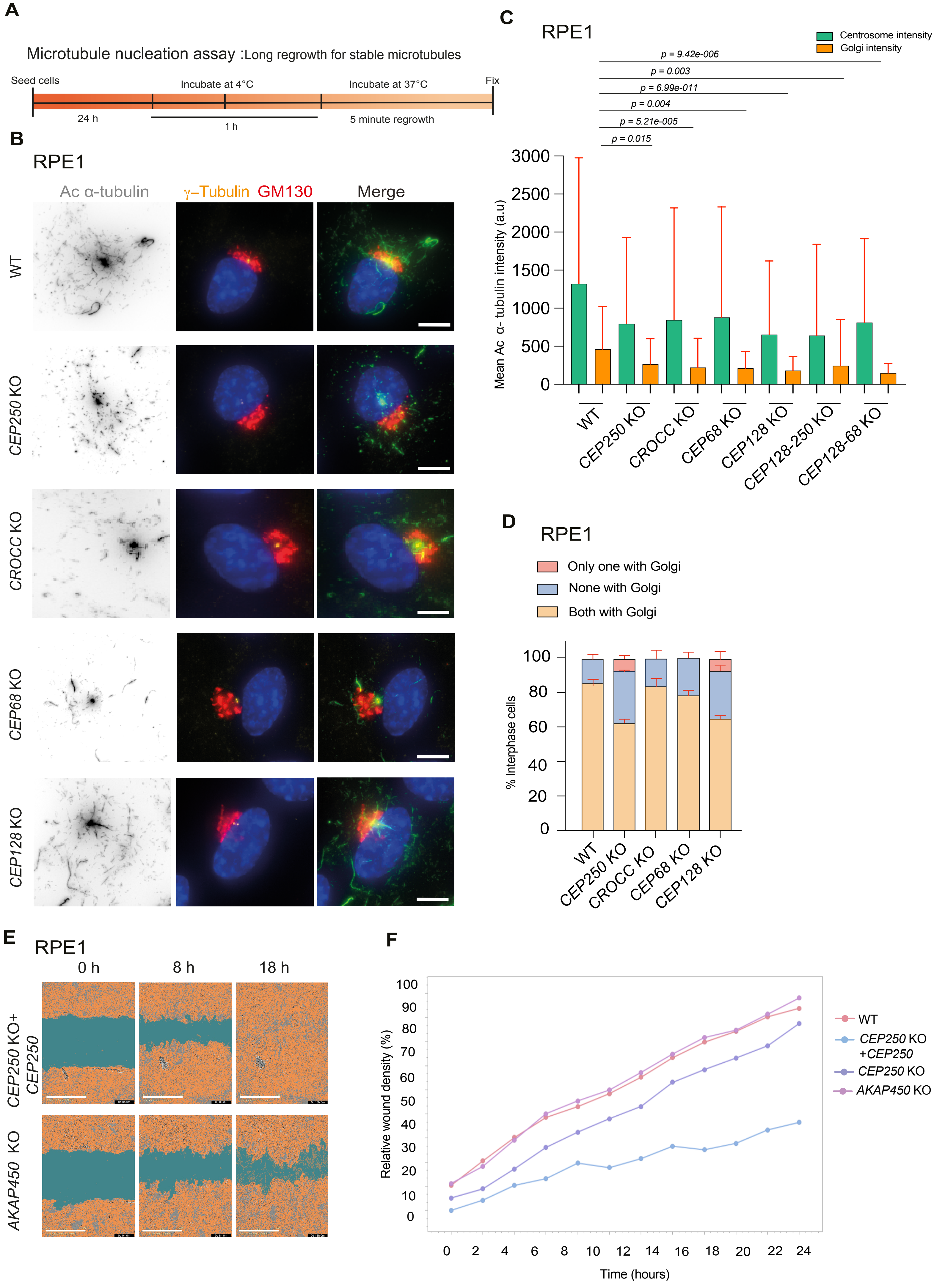
