## Appendix Figures S1-S8 for "Control of Golgi Architecture Through Three Centrosome Cohesion Pathways"

### Appendix Figure S1

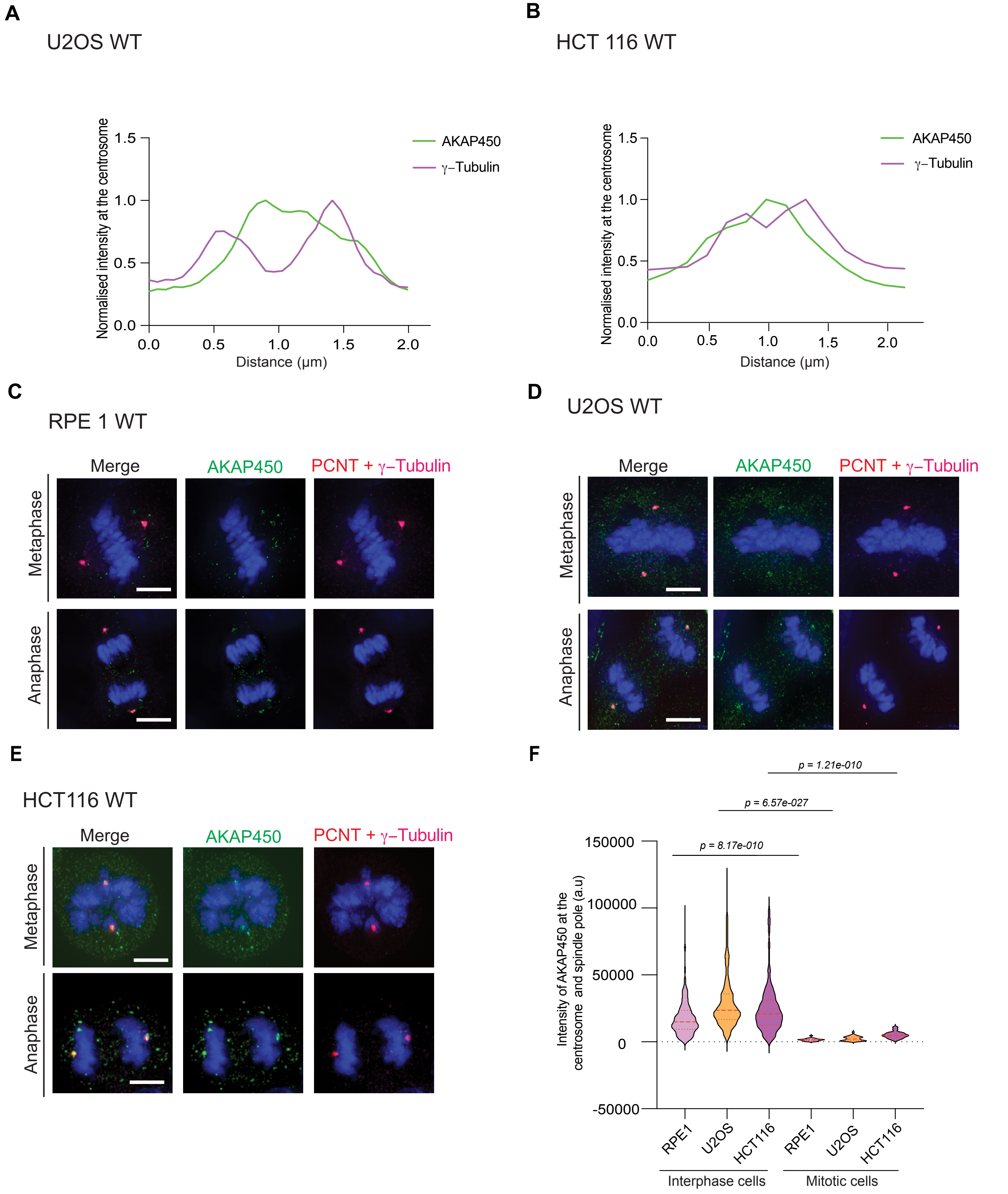

#### Appendix Figure S2

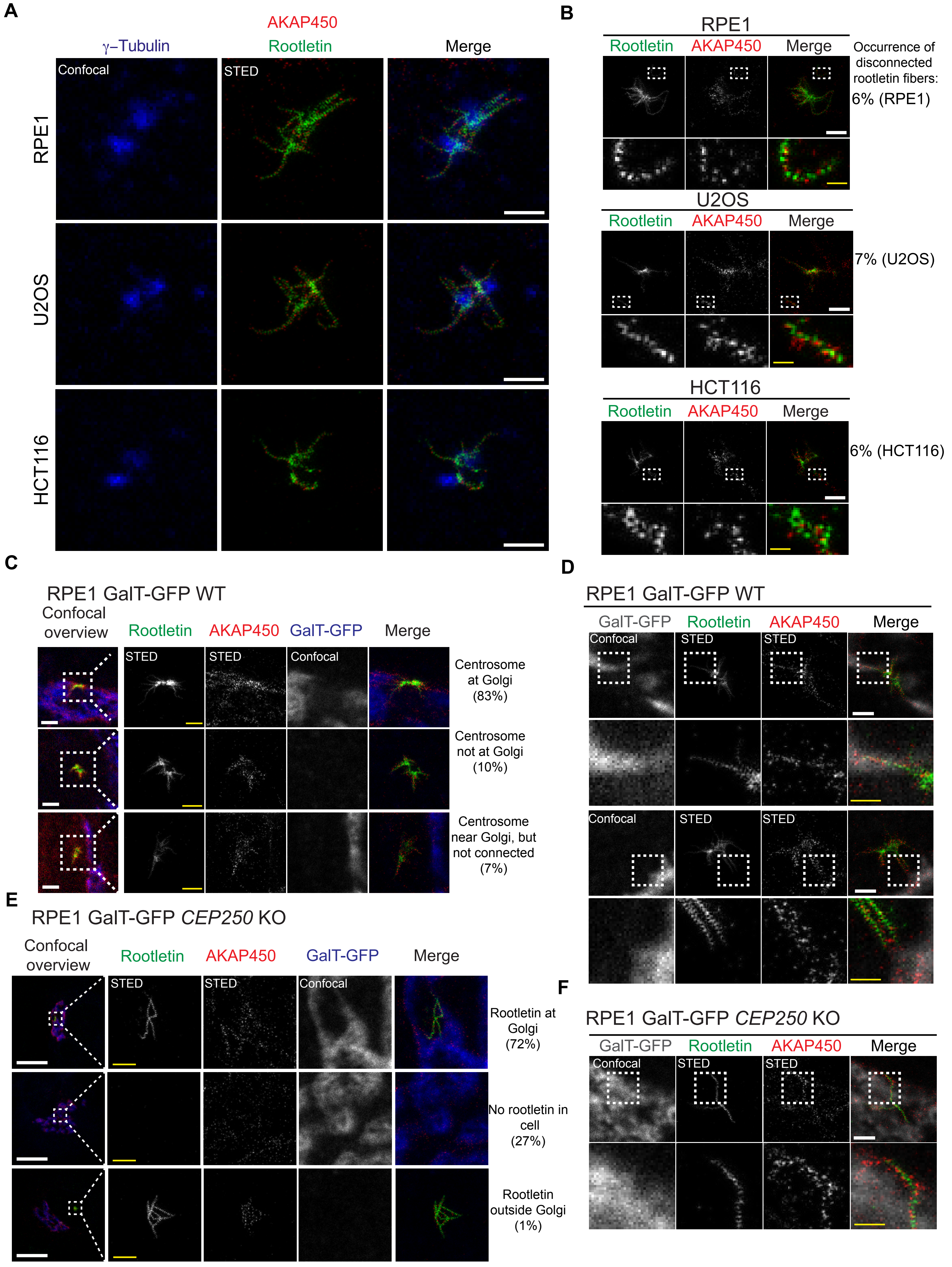

#### Appendix Figure S3

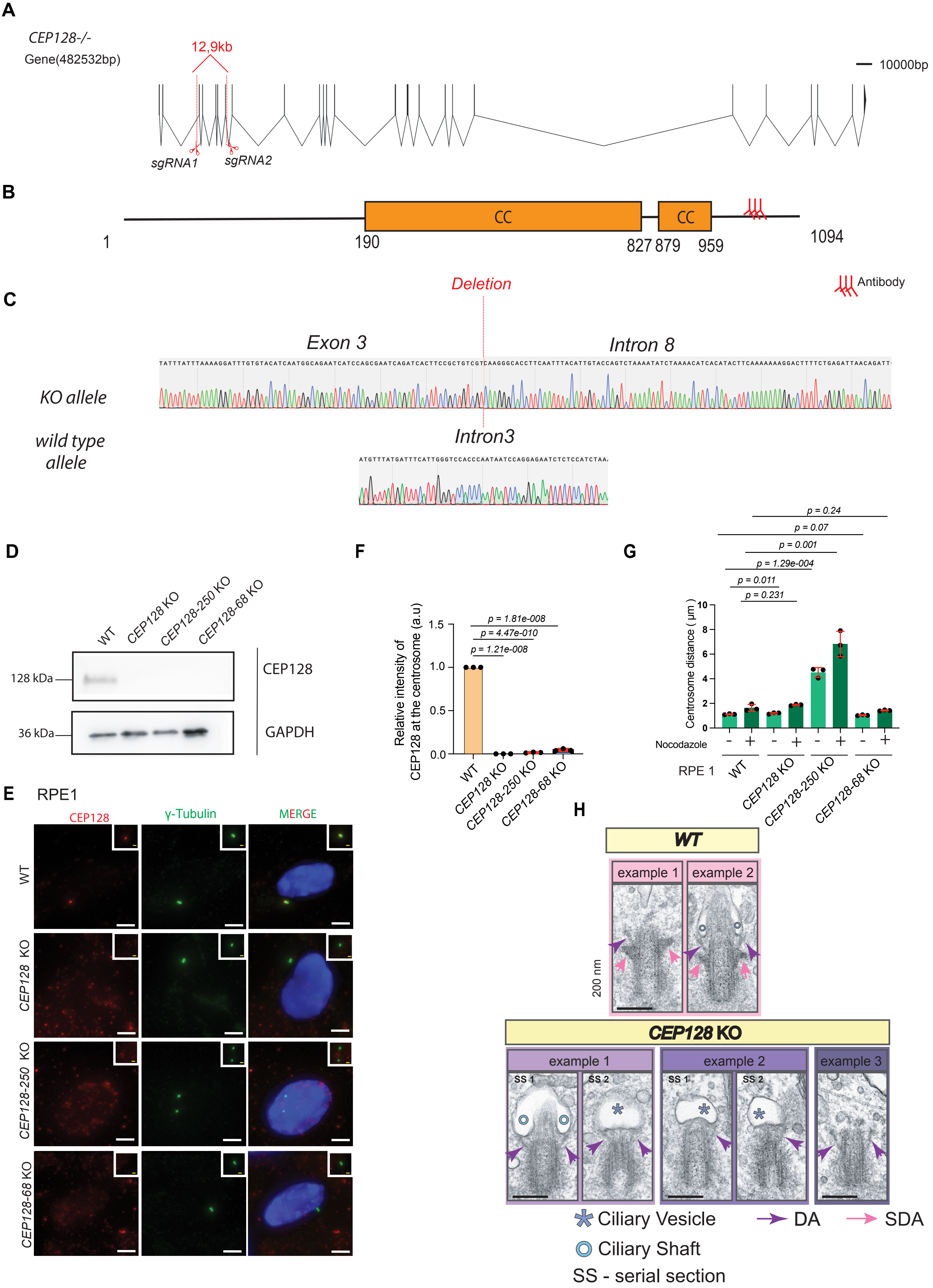

**Appendix Figure S4**

**A**

RPE1 *AKAP450$^{PACT}$-HA*

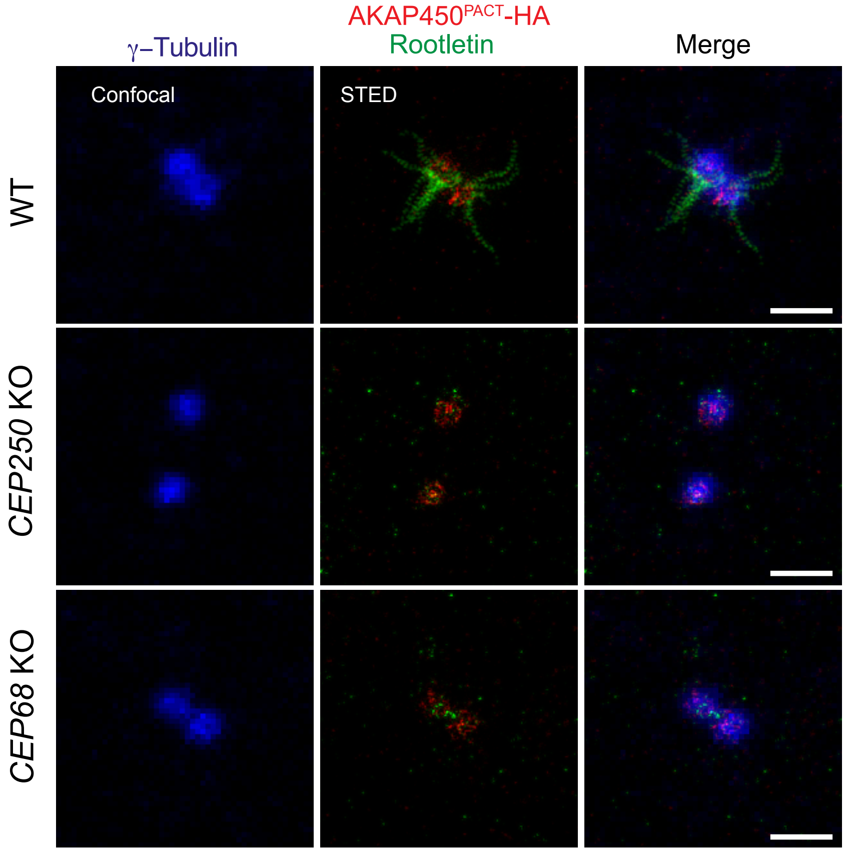


**B**

RPE1 *PCNT$^{PACT}$-HA*

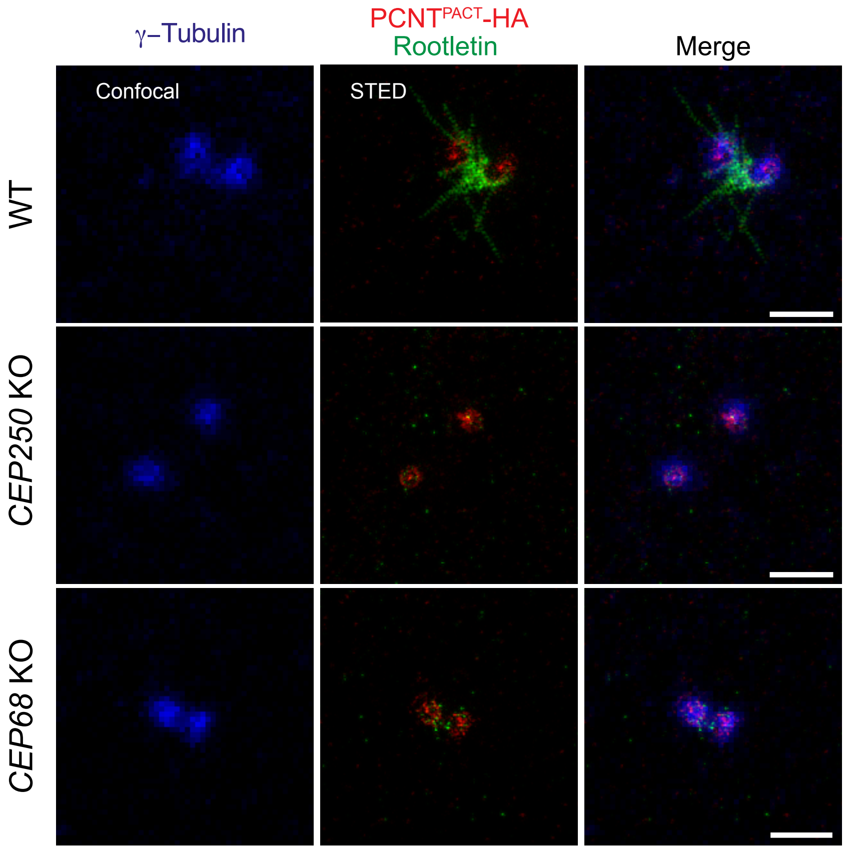

**Appendix Figure S5**

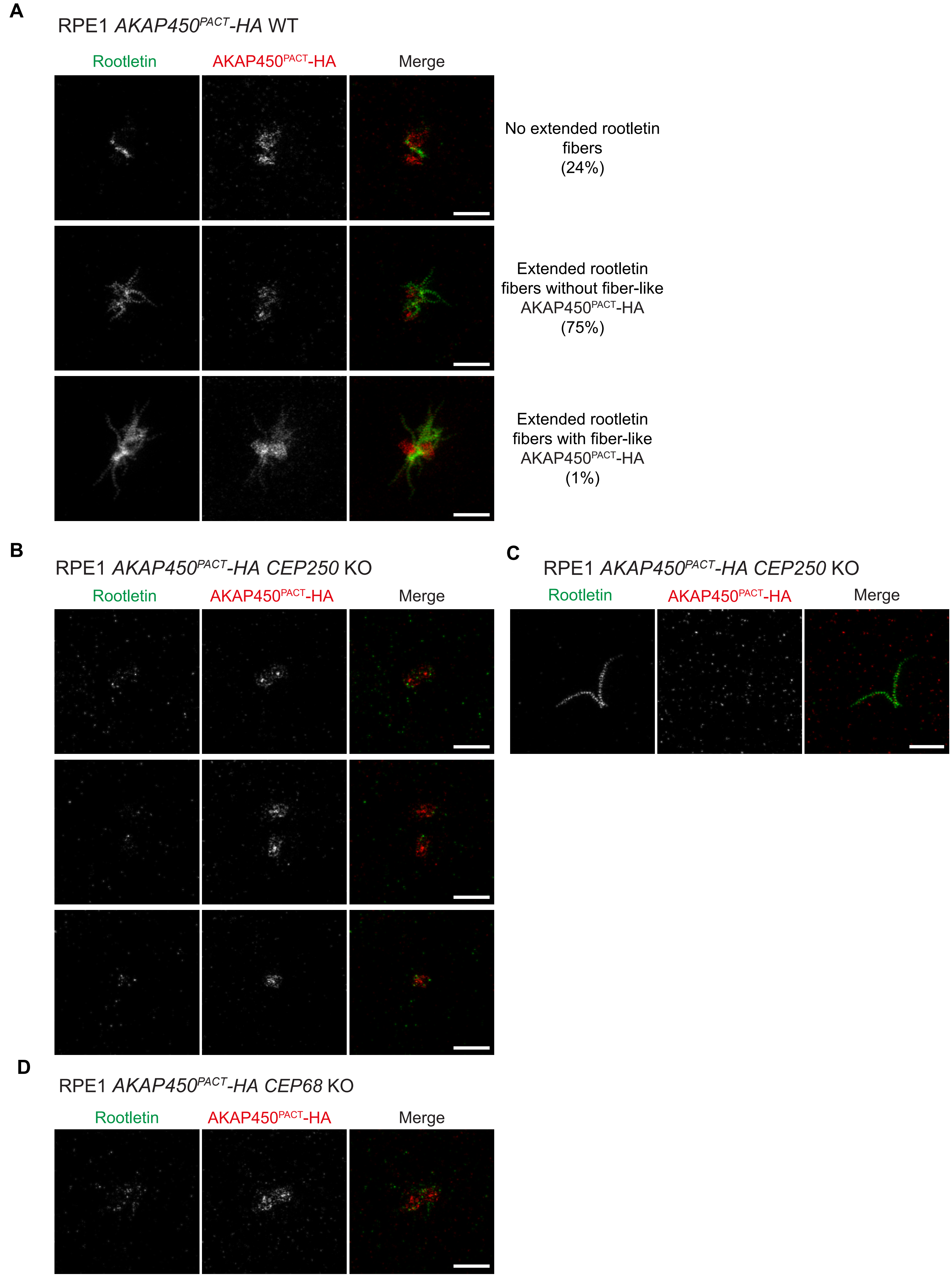

**Appendix Figure S6**

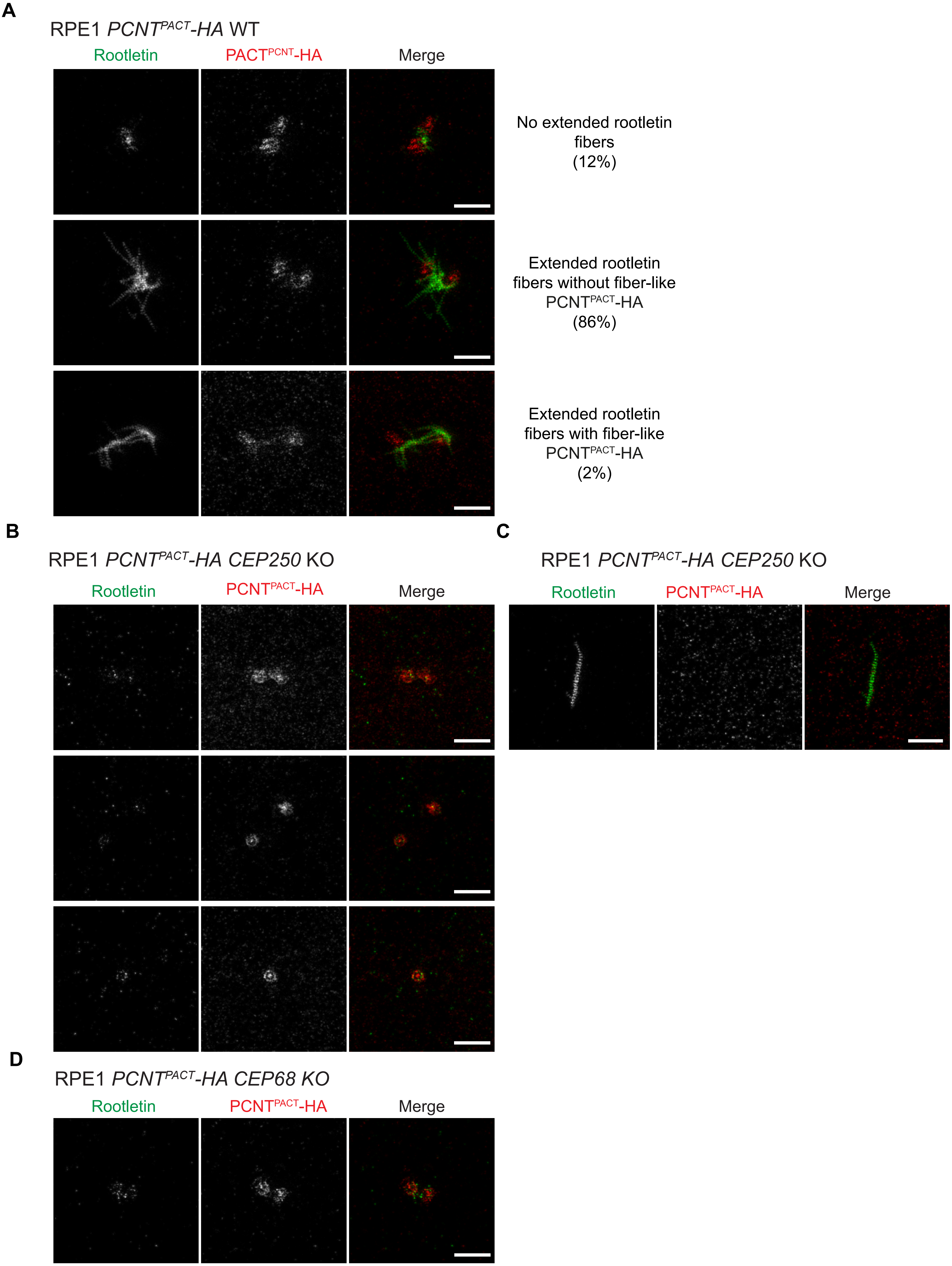

**Appendix Figure S7**

**A**

RPE1 WT

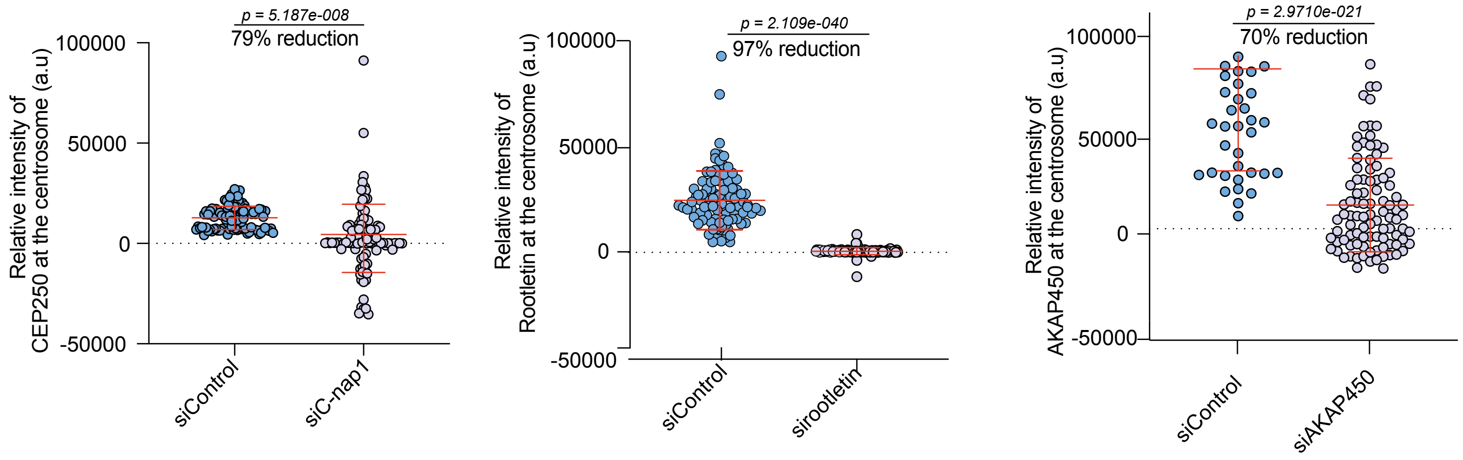


**B**

U2OS WT

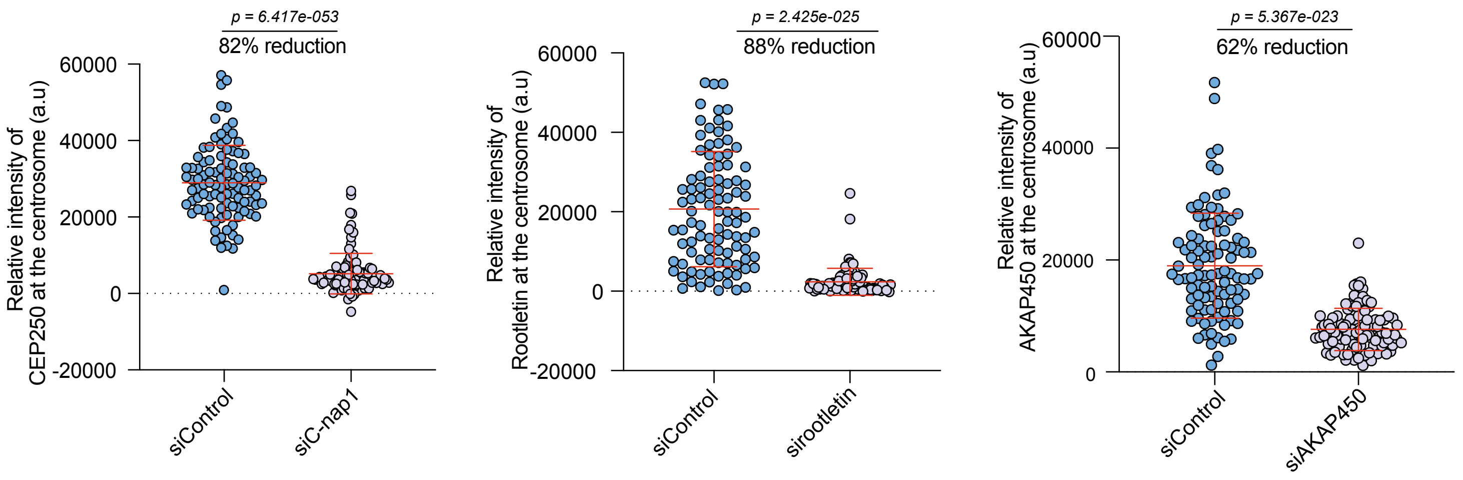


**C**

HCT116 WT

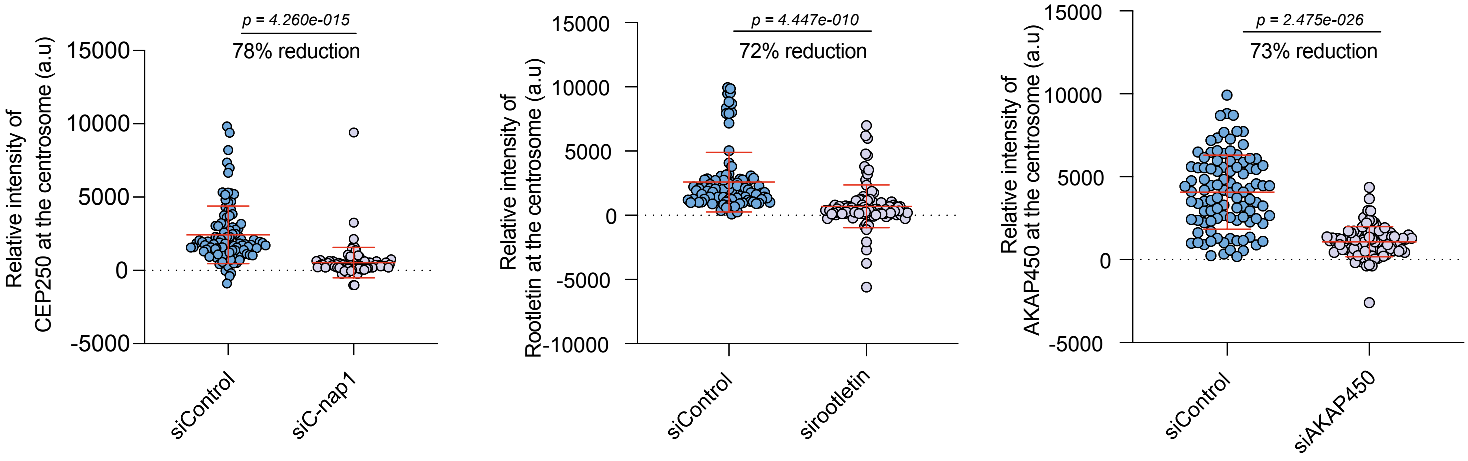

### Appendix Figure S8

**A**

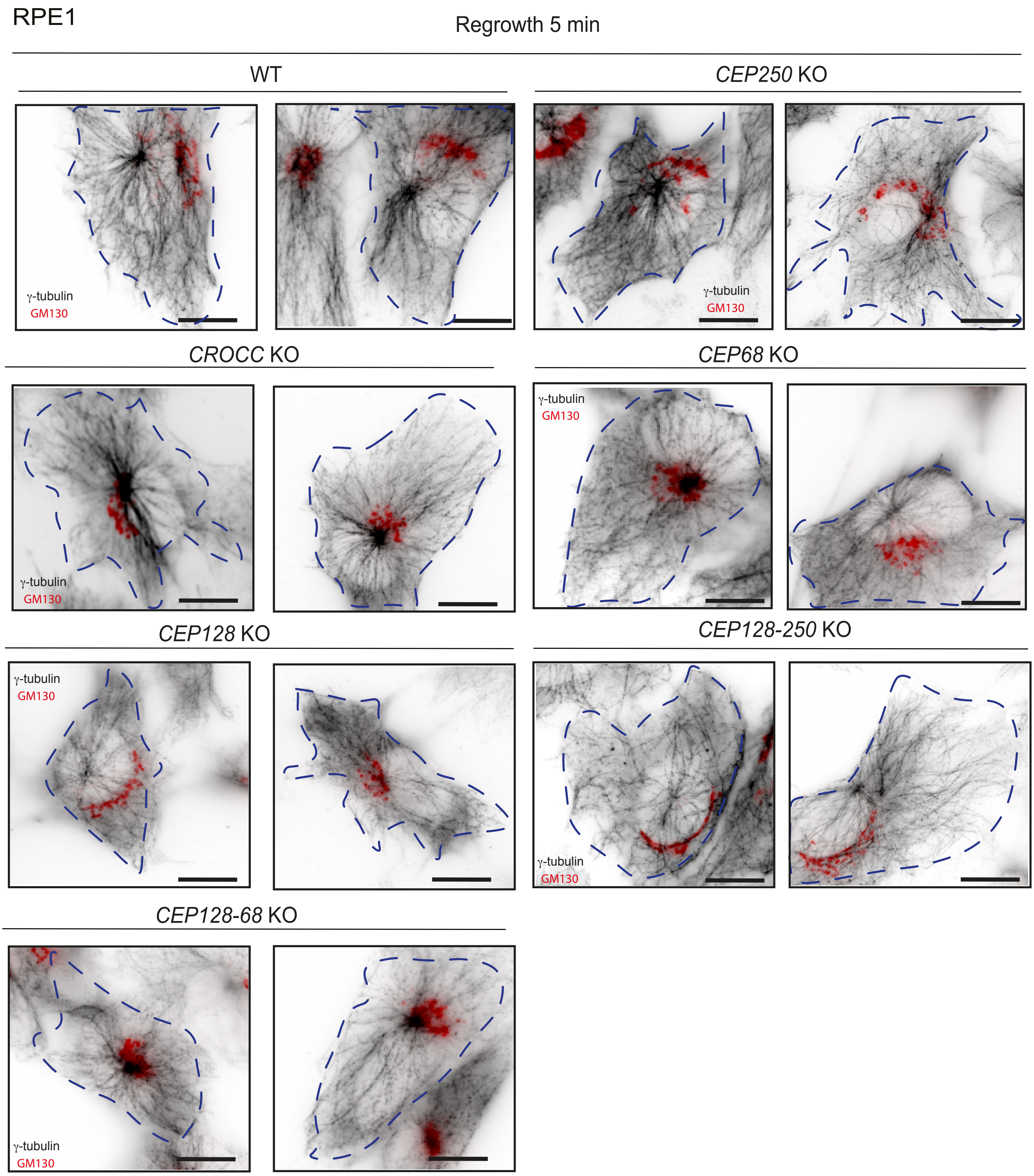


**B**

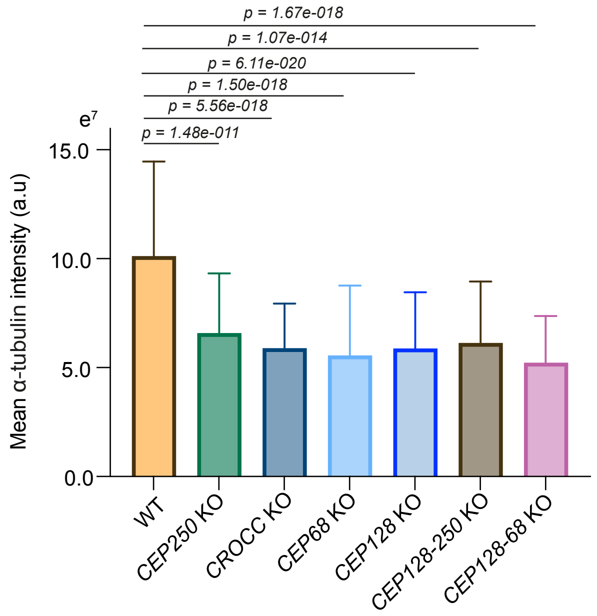
